# The heart rate discrimination task: a psychophysical method to estimate the accuracy and precision of interoceptive beliefs

**DOI:** 10.1101/2021.02.18.431871

**Authors:** Nicolas Legrand, Niia Nikolova, Camile Correa, Malthe Brændholt, Anna Stuckert, Nanna Kildahl, Melina Vejlø, Francesca Fardo, Micah Allen

**Affiliations:** Center of Functionally Integrative Neuroscience, Aarhus University Hospital, Denmark; Danish Pain Research Center, Aarhus University, Denmark; Aarhus Institute of Advanced Studies, Aarhus University, Denmark; Cambridge Psychiatry, University of Cambridge, United Kingdom

**Keywords:** heart rate discrimination, heartbeat tracking, interoception, psychophysics, metacognition

## Abstract

Interoception - the physiological sense of our inner bodies - has risen to the forefront of psychological and psychiatric research. Much of this research utilizes tasks that attempt to measure the ability to accurately detect cardiac signals. Unfortunately, these approaches are confounded by well-known issues limiting their validity and interpretation. At the core of this controversy is the role of subjective beliefs about the heart rate in confounding measures of interoceptive accuracy. Here, we recast these beliefs as an important part of the causal machinery of interoception, and offer a novel psychophysical “heart rate discrimination“ method to estimate their accuracy and precision. By applying this task in 223 healthy participants, we demonstrate that cardiac interoceptive beliefs are more biased, less precise, and are associated with poorer metacognitive insight relative to an exteroceptive control condition. Our task, provided as an open-source python package, offers a robust approach to quantifying cardiac beliefs.

**Highlights:**

- Current interoception tasks conflate cardiac beliefs with accuracy.
- We introduce a Bayesian method for estimating cardiac belief accuracy and precision.
- Individuals underestimate their heart rate by -7 BPM (95% CI [-8.6 -5.3]) on average.
- Cardiac beliefs are associated with reduced precision and metacognitive insight.
- The task and modelling tools are provided in the Python Cardioception Package.

## Introduction

Interoception denotes the ability to sense, perceive, and regulate internal visceral states (Chen et al., 2021; Sherrington, 1952). This ability is thought to depend on unique neurobiological pathways, which underpin the affective and somatic axes of selfhood (Craig, 2002; Critchley & Garfinkel, 2017; Seth & Tsakiris, 2018; Strigo & Craig, 2016). Measuring the individual interoceptive capacity to detect visceral signals, such as those arising from the lungs, heart, or stomach has recently come to the forefront of psychological and psychiatric research (Khalsa et al., 2018; Khalsa & Lapidus, 2016). A critical objective of this work is to determine the mechanisms by which interoception interacts with cognition and emotion, to ultimately derive sensitive and specific neuropsychiatric biomarkers from individual indices of visceral sensitivity. The majority of studies along these lines attempt to measure “interoceptive accuracy” (iACC) in the cardiac domain, as measured by the Heartbeat Counting (HBC) task (Dale & Anderson, 1978; Schandry, 1981), and similar heartbeat tracking or tapping tasks (Flynn & Clemens, 1988). While easy to implement, these tasks suffer from serious methodological challenges that obscure their interpretation. To overcome these challenges, we developed a novel psychophysical approach to measure the accuracy, bias, and precision of interoceptive beliefs in the cardiac domain.

The measurement of interoceptive accuracy presents a unique challenge compared to that of exteroception: unlike vision or touch, the heart is not typically amenable to direct experimental control. The inability to control the information present in the stimulus (e.g., the heartbeat) places hard constraints on interoception research, such that most extant tasks ask participants to count uncontrolled endogenous states (e.g., heartbeats) or to determine whether exteroceptive stimuli are synchronized with said states. While these tasks are widely used, they suffer from several confounds which place strong limitations on their reliability, interpretability, and validity (for review see Brener & Ring, 2016; Desmedt et al., 2018; Desmedt, Corneille, et al., 2020; Ring & Brener, 2018; Zamariola et al., 2018).

A central issue associated with the use of the HBC or similar tasks concerns the role of subjective beliefs about one’s heart rate. These simple measures require participants to silently attend to and count their heartbeats for various intervals, or to tap in rhythm to felt beats. Several authors point out that participants could exploit various strategies to increase their accuracy (Clemens, 1979; Flynn & Clemens, 1988; Pennebaker & Hoover, 1984). Crucially, even when the heart rate is directly modulated by as much as 60 beats per minute (BPM) via pacemaker, counted heartbeats showed little alteration beyond expectations about different sitting or standing postures on the heart rate (Windmann et al., 1999). Accordingly, participants’ subjective prior beliefs about the heart rate have been repeatedly found to be more predictive of counts than actual heartbeats (Ring & Brener, 1996), and it has been shown that these beliefs can be manipulated via false feedback independently of any true change in heart rate (Ring et al., 2015). A more accurate prior knowledge about one’s heart rate, e.g. amongst medical practitioners or athletes, can influence HBC accuracy scores (Murphy et al., 2018), such that when explicitly instructing participants not to estimate beats, but to instead count felt ones, this bias is reduced (Desmedt et al., 2018). More recently, the validity of the HBC task has been further questioned by reports showing that interoceptive accuracy scores are largely driven by under-counting (Zamariola et al., 2018), suggesting that HBC-derived scores are merely a rough reflection of subjective beliefs about the heart rate (Desmedt, Luminet, et al., 2020).

These reports raise serious concerns given the rising interest in interoceptive measurements as potential psychiatric biomarkers (Eggart et al., 2019; Forkmann et al., 2019; Paulus & Stein, 2010). This poor construct validity could also explain why little to no relationship between HBC-derived scores and various psychiatric symptom measures has been found at the meta-analytic level (Desmedt, Houte, et al., 2020). Here, we argue that the inconsistencies between these HBC-derived scores and interoceptive ability could be better handled by more rigorous measurement and modelling of the role of subjective beliefs in cardiac interoception. Although these tasks were originally designed to be objective and selective measures of the ability to detect afferent cardiac sensory information, they fail to account for factors confounding score variances, such as prior beliefs about the heart rate and other common introspective or self-report biases. In particular, these approaches struggle to dissociate interoceptive sensitivity, bias, and accuracy, confounding the role of subjective vs. objective performance in interoceptive measures.

Another commonly used task, the Heartbeat Discrimination (HBD) task (Whitehead et al., 1977), suffers from different, but similarly serious drawbacks. This method presents participants with a series of tones whose onset times are delayed at different intervals relative to the R-wave. Tones presented approximately at systole (typically, R + 170 ms) are treated in signal theoretic terms as the “signal plus”, while tones presented at a variable time after systole (typically, R + 300 ms) are treated as “signal minus”. This design is based on strong assumptions about when, relative to the cardiac phase, participants are most likely to feel the heartbeat. These assumptions have been challenged by results obtained using a similar task based on a method of constant stimuli (MCS), where tones are presented at 5 different intervals with respect to the R-wave.

Using this MCS-based method, Brener and colleagues demonstrated that individuals vary substantially in terms of when relative to r-wave, heartbeats are perceived (Brener et al., 1993; Brener Jasper & Ring Christopher, 2016; for review see Ring & Brener, 2018) and that calibrating the HBD offset intervals to each subject improves detection scores to above chance, seriously undermining the notion that the HBD can be used as a signal theoretic measure to delineate cardiac sensitivity and bias. However, while the MCS likely improves the quantification of single-beat detection when compared to the HBD, both tasks require participants simultaneously to attend to exteroceptive and interoceptive multi-sensory inputs, a difficult cognitive task that further obscures the relationship to interoception. Moreover, the MCS requires long testing times (as much as 1 hour), which can be problematic for clinical populations, and further requires sophisticated equipment capable of achieving precise cardiac-tone synchrony and stimulus time. As such, there is a need for robust, accessible measures that are amenable to clinical settings, and which can flexibly dissociate the bias, sensitivity, and precision of cardioceptive decisions.

To achieve this, we developed the heart rate discrimination task (HRD), a novel psychophysical approach to quantifying cardioceptive decisions. Through Bayesian modelling of cardiac psychophysics, the HRD delineates the accuracy of cardiac beliefs into the bias (i.e., the error between the perceived HR versus ground truth), and precision (i.e., the uncertainty around this estimate) of trial by trial cardiac decisions. By presenting stimuli dynamically across trials, titrated to the current heart rate, this approach estimates psychometric perceptual and metacognitive curves indicating participants’ ability to update and monitor cardiac beliefs under different conditional manipulations.

To demonstrate the utility of the HRD for measuring cardiac beliefs, characterize the overall interoceptive psychometric function, and establish the face validity of this approach, we measured HRD performance in 223 participants at a resting heart rate while seated upright in a standard testing booth, together with heartbeat counting scores. To further quantify internal (test-retest) reliability, we re-tested HRD performance in the same participants following 6 weeks. Our results demonstrate that cardioceptive beliefs are reliably and robustly measured by the task across both sessions. Further, we find that cardioceptive beliefs are more negatively biased, imprecise, and associated with poorer metacognitive insight relative to an exteroceptive control condition.

## Methods

### Heart Rate Discrimination task

#### Task Overview

To measure the bias, precision, and metacognitive calibration of interoceptive beliefs, we created the Heart Rate Discrimination task (HRD) (See **Fig. 1**). The goal of the HRD is to provide an efficient method for measuring the influence of subjective beliefs, viscerosensory inputs, and other possible contextual factors on interoceptive decisions about heart rate. The task asks participants to first attend to their cardiac sensations and then to decide on each trial whether a “feedback” tone series is faster or slower than their heart rate in a 2-Interval Forced Choice design (2-IFC). To control for possible non-interoceptive processes such as working memory or general temporal estimation biases, we also implemented an exteroceptive control condition, in which participants had to discriminate whether a series of tones was faster or slower than another “reference” sequence of tones.

**Legend Figure 1:**
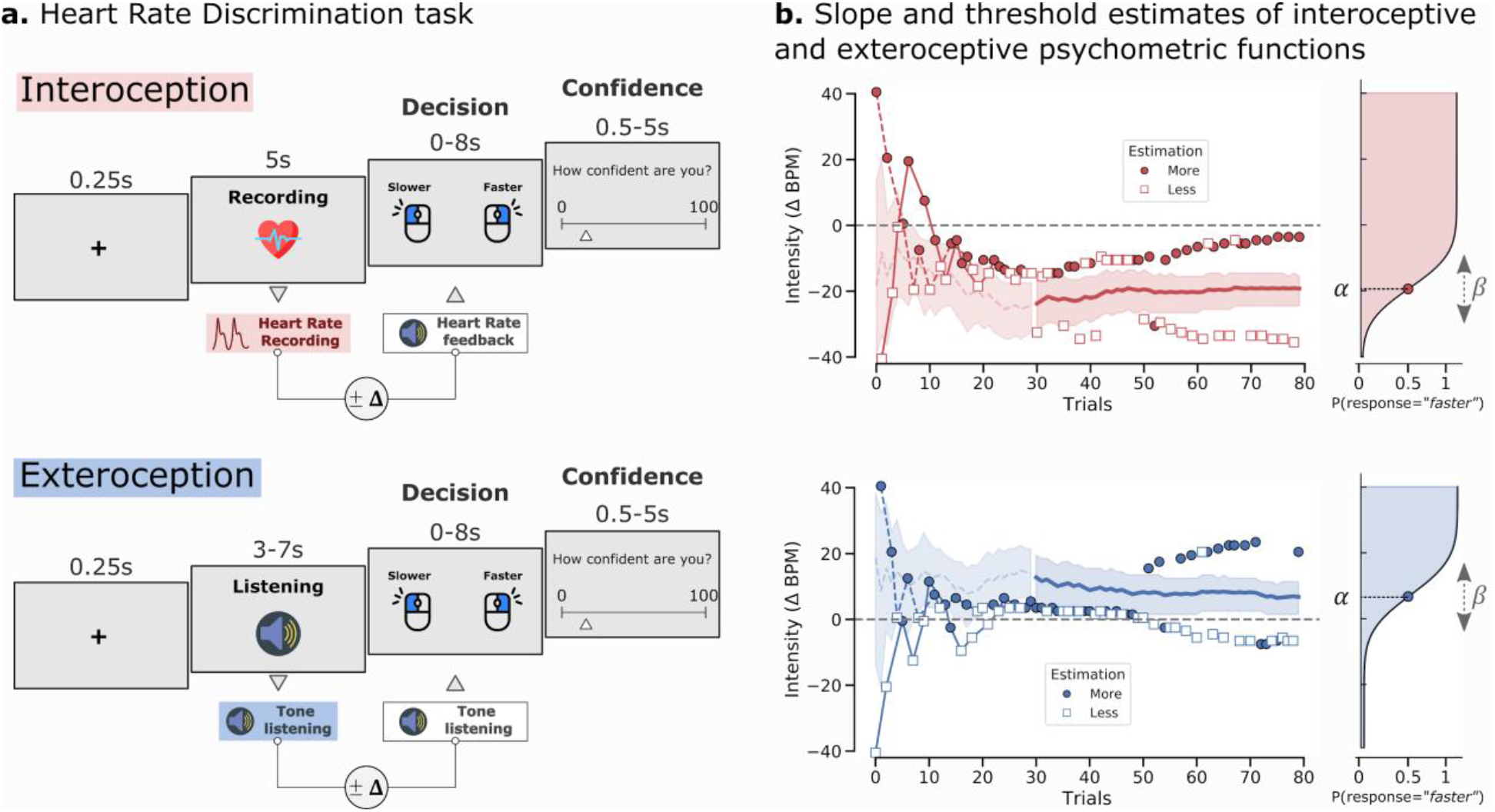
**A. Heart Rate Discrimination Trial Design (Session 1).** Participants were presented with 160 trials testing their exteroceptive (blue) and interoceptive (red) bias and precision (80 in each condition in randomised interleaved order). During interoceptive trials, participants were instructed to attend to their heart rate for 5 seconds, while it was recorded using a pulse oximeter. The average heart rate for the trial was then computed and used to select the frequency of the tones presented during the decision phase, increased or decreased by an intensity value generated by the staircase, i.e. **Δ**-BPM. During exteroceptive trials, a sequence of tones was presented to the participant with a frequency between 40 and 100 BPM, drawn randomly from a uniform distribution. This value then determined the frequency of the tones presented during the decision phase, increased or decreased by a value generated by the staircase procedure. **Δ**-BPM values were controlled by separate staircases for each condition. To estimate metacognitive ability for each modality, at the end of each trial, participants were asked to rate their subjective decision confidence (from 0 - guess to 100 - certain). **B. Staircases for each condition from an exemplary subject.** Trials classified by participants as faster or slower are depicted with circles or squares respectively. The shaded area represents the 95% CI of the threshold posterior distribution. On the right panel, the resulting cumulative normal distribution is plotted using the final parameters estimated by the Psi procedure.

To estimate psychometric functions for both conditions, we applied a well-established adaptive Bayesian psychophysical method (“Psi”) (Kingdom & Prins, 2016; Kontsevich & Tyler, 1999; Prins & Kingdom, 2018). This technique adaptively estimates the probability of a participant responding that the feedback tones were “faster” or “slower” than the true heart rate (interoception), or the reference tone (exteroception) on each trial, given the frequency difference between the two stimuli, or **Δ**-Beats Per Minute (**Δ**-BPM). This procedure estimates the point of subjective equality (PSE) both for interoceptive and exteroceptive decision processes in the same **Δ**-BPM units. The PSE is henceforth referred to as the threshold of the psychometric function. This threshold represents the difference between the true frequency of the heart rate and the estimated cardiac frequency by the participant (see **Fig. 1.** the threshold is denoted **α**). A negative threshold, therefore, indicates the degree to which a participant underestimates their cardiac frequency, while a positive threshold indicates an overestimation. In addition to the threshold measure, the procedure further estimates the slope of the psychometric function (denoted **β** in **Fig. 1.**), which represents the precision, or uncertainty, around this estimated perceptual bias, also in units of **Δ**-BPM. A larger slope value reflects a less steep psychometric curve, indicating increased uncertainty (i.e. reduced precision) in the cardioceptive decision process.

#### HRD Trial Design

During interoceptive trials, participants silently attend to their heart rate for 5 seconds, e.g., in a “heart listening” phase (see **Fig.1**), during which the heart rate is monitored using a soft-clip pulse oximeter placed on one of the fingers of the non-dominant hand. The raw signal is analyzed in real-time using a systolic peak detection algorithm, and the heart rate is calculated as the average of the inter-pulse interval (see **Physiological Analyses**, below). After this listening phase another sequence of five auditory tones was presented (frequency: 440 Hz; duration: 200 ms). This “two-interval” design is a deliberate choice so that participants attend solely to their cardiac sensation before the presentation of auditory feedback.

At any point during the auditory feedback, the participant can press the right or left mouse button to indicate whether the feedback sequence was faster or slower than their estimated average heart rate, terminating the response and feedback period. The maximum response time for the type 1 decision task was 8 seconds. Following the decision interval, participants provide a subject confidence rating (from 0 - uncertain to 100 - certain, minimum possible response time: 0.5 seconds; maximum: 5 seconds), and the next trial begins. To prevent motor preparation of the confidence rating, the starting point of the rating scale cursor is randomly jittered around the midpoint by about +/- 70% of the scale length.

Crucially, the frequency of the second tones was adjusted to the frequency of the first tones (exteroceptive modality) or the recorded cardiac frequency (interoceptive modality). This difference is denoted **Δ**-BPM and corresponds to the stimulus intensity manipulated by the staircases (see the *Staircase procedure* section below). For example, if the heart rate recorded during the listening condition is 60 BPM, and the **Δ**-BPM value is −15, the feedback tone frequency will be set to 45 BPM. In this example, if the participant answers “Slower”, this is considered a correct answer, otherwise, this is considered an incorrect answer. In this way, the staircase procedure hones in on the point of subjective equality, or threshold (**α**), at which the participant is equally likely to respond “Faster” or “Slower”.

During exteroceptive trials, participants compared two sequences of tones, instead of comparing their heart rate with the feedback tone sequence. Here, the first (“reference”) tone sequence frequency was randomly selected from a uniform distribution (lower bound = 40 BPM, upper bound = 100 BPM, signal frequency: 440 Hz; tone duration: 200 ms), and the second tone sequence frequency presented at a BPM above or below this value as determined by the staircase procedure. After this listening phase, the participants underwent the same decision and confidence task as in the interoceptive trials, that is, to decide whether the second sequence was faster or slower than the first one. The tone presentation ceased when the response was provided (maximum response time: 8 seconds). As in the interoceptive condition, the intensity **Δ**-BPM was adjusted across trials using the same adaptive Bayesian approach for estimating threshold and slope.

#### Adaptive Staircase Procedure

The primary aim of the HRD task is to estimate the difference between the objective heart rate and the participant’s subjective perception of this heart rate (i.e., the threshold α), as well as the precision or uncertainty around this perceptual belief (i.e., the slope **β**). The HRD achieves this via an adaptive staircase procedure which manipulates the **Δ** value that was added to the true BPM to produce the feedback tone frequency. This psychophysical procedure can be described as an appearance-based 2-Interval Forced Choice (2-IFC) similar to the 2-Alternative Forced Choice procedure implemented in the Vernier-alignment task, (Kingdom & Prins, 2016, Chapter 3.3), where varying the degree of difference between two stimuli allows estimating the threshold (**α**) and the slope (**β**) of the underlying decision process. In our implementation, HRD thresholds are adaptively estimated using either two interleaved standard n-up/n-down staircases (the first steps were manually fixed to: {20, 12, 12, 7, 4, 3, 2, 1}; starting value: −40.5 and 40.5) (Dixon & Mood, 1948), or using an adaptive method known as Psi (Kontsevich & Tyler, 1999).

Psi is a Bayesian adaptive psychophysical method that manipulates the **Δ**-BPM deviation values and estimates the slope and the threshold of the underlying sensory psychometric function (Kingdom & Prins, 2016; Prins & Kingdom, 2018). The psychometric function relates the deviation **Δ** to the proportion of “Faster” decisions using the following formula:

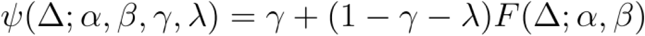

in which **Δ** refers to stimulus intensity, ***ψ*** refers to the proportion of trials rated as faster by the participant, and **γ** and 1-**λ** are nuisance parameters corresponding to the lower and upper asymptote, respectively. ***F*** is the cumulative normal distribution parameterized using threshold **α** and slope **β**. The parameter **λ** is often referred to as the *lapse rate* and describes the probability of a stimulus-independent negative response (here, the probability of answering “slowed” regardless of the frequency of the tones). In appearance-based 2-IFC tasks like the HRD, the lapse rate determines both **γ** and 1-**λ**. Here, this parameter was fixed, because it is assumed that responses obtained from this kind of task only contain limited information on the real value of this parameter. Further, the estimation of the lapse rate as a free parameter can potentially introduce bias (Prins, 2012). Here, we assumed that **γ**=**λ**=0.02.

Irrespective of the staircase procedure (i.e., n-up/n-down or Psi), it should be noted that the psychometric function was not fitted to the proportion of correct trials given the **Δ** intensity, but on the probability of a participant making a “Faster” response given the **Δ** intensity (see **Fig. 1b**). This procedure is known as an appearance-based staircase and entails that the probability of a participant answering “Faster” given an increasing **Δ**-BPM value is expected to follow a monotonic psychometric function. However, this is not the case for the probability that the response is correct relative to the ground truth HR. Here, the probability of answering correctly increases with **Δ**-BPM increments towards either positive or negative infinity and is 0.5 around the threshold. As a consequence, the online estimation of an accuracy-based staircase requires dedicated adaptive methods for non-monotonic psychometric functions (García-Pérez, 2014).

### Python Cardioception Package

We implemented two interoception tasks using Psychopy v3.2.3 (Peirce et al., 2019), the classic Schandry Heartbeat Counting Task (Dale & Anderson, 1978; Schandry, 1981), and the Heart Rate Discrimination task. The code for the two interoception tasks is made publicly available in the Cardioception Python Package (https://github.com/embodied-computation-group/Cardioception). The package natively supports recording and processing pulse rate activity as recorded by the Nonin 3012LP Xpod USB pulse oximeter together with a Nonin 8000SM “soft-clip” fingertip sensor (https://www.nonin.com/) by interfacing with the Systole python package for pulse oximetry (Legrand & Allen, 2021).

### Participants

223 participants between the ages 18 and 56 (130 females, 93 males, 1 other, age = 25.0 ± 5.50) participated in the study. They were recruited through the Center of Functionally Integrative Neuroscience (CFIN) SONA system participant pool, local advertisements and flyers, social media, and the aarhusbrain.org website. All measures took place at Aarhus University Hospital, Denmark, and were performed on a computer in a behavioural testing room. All participants had normal or corrected to normal vision, and were of at least average proficiency in both Danish and English. All participants were healthy and did not take psychoactive, psychiatric, or cardiovascular medications. Participants who could potentially be pregnant or were breastfeeding had MRI contraindications (e.g., claustrophobia), or reported that they were not able to abstain from alcohol and drugs 48 hours before study participation, were not included in the study. All participants took part in a larger experiment including multiple brain scans, psychiatric inventories, and other behavioural measures (data not reported here). Participants were invited to complete two separate experimental sessions of the HRD task (henceforth referred to as “Session 1” and “Session 2”), separated by 46.89 days on average (min=10; max=97; std=23.87, statistics from participants that completed both sessions). Among the 223 participants, 218 participants completed session 1 and data were lost for 5 of them due to technical difficulties. 192 participants (86% of the total sample) completed session 2. 11 participants were excluded from session 1 due to poor signal quality and/or for not having performed the HRD task correctly (detail concerning the exclusion of participants based on insufficient staircase convergence are provided in Supplementary Material). 1 participant was excluded from session 1 and 2 due to a prior psychiatric diagnosis. After exclusion, 206 participants had completed Session 1 and 191 participants had completed Session 2. Among the 191 participants that completed session 2, 179 had also completed session 1 (5 were removed due to technical difficulties, and 7 were removed due to signal quality). When analysing confidence ratings, 1 participant from Session 1 and 1 participant from session 2 were dropped due to insufficient variance for estimation of M-ratio (see **Analysis**). 214 participants completed the HBC task, and 7 of them were removed due to poor physiological signal quality. 193 participants had completed both the HBC task and the HRD task during session 1. Participants received monetary compensation for each session (350 DKK). The study was conducted in accordance with the Declaration of Helsinki and was approved by the Region Midtjylland Ethics Committee.

### Physiological recordings

Physiological signals in both sessions were recorded using the Nonin 3012LP Xpod USB pulse oximeter together with a Nonin 8000SM “soft-clip” fingertip sensor (https://www.nonin.com/) by interfacing with the “Systole” python package (v0.1.3) for pulse oximetry (Legrand & Allen, 2021). Previous work reported that the pressure exerted by some pulse oximeters on the surface of the skin could provide sensory feedback to some participants, therefore potentially biasing the estimation of interoceptive accuracy (Murphy et al., 2019). We selected the Nonin Pulse softouch pulse oximeter as Murphy and colleagues demonstrated that this device did not elicit fingertip pulse sensations. We further attempted to mitigate this effect by asking the participants to move the device from the index finger to another fingertip if they felt any such sensory feedback, although we did not record the proportion of participants for which this adjustment was made. Participants were further asked to keep their hand still on the table or on their thighs so as to not introduce heart rate measurement errors, due to movements.

#### Heartbeat Counting Task

The Heartbeat Counting (HBC) task is perhaps the most widely used and easily implemented task for measuring interoceptive accuracy (Dale & Anderson, 1978; Schandry, 1981). However, this procedure has been previously criticized, in part because it could be confounded by beliefs about one’s heart rate (Desmedt, Corneille, et al., 2020). To better validate and interpret our new HRD measure, we implemented a revised version of the HBC (Garfinkel et al., 2015), together with specialized instructions to reduce the role of bias in the derived interoceptive accuracy (iACC) scores. Considering the substantial evidence that HBC scores are influenced by beliefs in the heart rate, we expected to observe significant correlations between HBC iACC scores and individual HRD thresholds. Participants were asked to count their heartbeats for various periods of time while sitting silently. The HBC task consisted of 6 trials and lasted 25, 30, 35, 40, 45 or 50 seconds. The order of the trials was randomized across participants. The HBC task was measured only in Session 1.

#### HRD Task Procedure - Session 1

At Session 1, the task comprised 160 trials, equally distributed between the interoceptive and the exteroceptive conditions. For each condition, the first 30 trials were run using an adaptive 1-up/1-down staircase procedure, and the remaining 50 trials were run using a Psi procedure (Kontsevich & Tyler, 1999). Our intention in combining these two staircase procedures was to ensure that each experiment started with a mixture of trials that would be clearly perceived as “Faster” or “Slower” by the participant. The initial up/down staircases consisted of 2 randomly interleaved 1-up 1-down staircases per condition initialized at high (**Δ**-BPM = 40) and low (**Δ**-BPM = −40) starting values following the recommendations of Cornsweet (Cornsweet, 1962). The Psi staircases were initialized at starting values informed by the 1-up 1-down staircases, achieved by updating Psi in the background with the intensity values and responses recorded during the first 30 up/down trials. Based on our pilot studies, Psi was initialized such that the prior for **α** was uniformly distributed between −40.5 and 40.5 and the prior for **β** was uniformly distributed between 0.1 and 20. The **α** precision was 1 BPM to ensure that the intensity **Δ**-BPM = 0 was excluded a priori and never presented. The **β** precision was set to 0.1.

The main task was preceded by a tutorial phase, which comprised 5 interoception and 5 exteroception trials with accuracy feedback after the decision and without confidence ratings, as well as 5 interoception trials without feedback, but with confidence ratings, as in the main experiment. For these trials, we fixed the absolute **Δ** value to 20 BPM and randomly selected for negative or positive differences at each trial. This was intended to clarify the instructions, to provide the participant with an easier version of the task, and ensure that they had an opportunity to practice and adapt before the staircase procedure, which can be biased by initial lapses. All auditory tones in Session 1 were presented through the stimulus PC speakers. The total duration of the HRD task was 31.31 minutes on average (*SD* = 3.32, *MIN* = 24.27, *MAX* = 42.46).

#### HRD Task Procedure - Session 2

To assess the internal (test-retest) reliability of HRD performance, all participants were invited back for a second testing session. Here, all aspects of the HRD were as in Session 1, minus the following, detailed below.

The total duration of the HRD at Session 2 was 22.69 minutes on average (*SD* = 2.45, *MIN* = 18.81, *MAX* = 31.81). Due to a change in our testing environment, which exposed participants to additional MRI noise, we opted to deliver auditory stimuli via over-the-ear headphones to limit external distractions. We also decreased the maximum decision time from 8 to 5 seconds. To optimize psychometric estimation, we made the following changes to the parameters and overall adaptive procedure. In particular, we observed a ceiling effect in the Session 1 slope parameters (See Supplementary **Fig. 2.b**), likely induced by an overly restrictive range on the slope prior distribution. To improve the estimation of this parameter, we increased this range from 0.1-20 in Session 1, to 0.1-25 in Session 2. The range of the threshold was increased from [−40.5, 40.5] in Session 1 to [−50.5, 50.5] in Session 2. We also simplified the staircase procedure in Session 2, running only the Psi staircase instead of the dual staircase approach described earlier. As the 1-up/1-down staircase initialization was intended to ensure participants heard a sufficient number of positive and negative **Δ**-BPM trials (i.e., trials in which the feedback was truly faster or slower than their true heart rate), we instead implemented “catch” trials presented at fixed intervals above and below zero **Δ**-BPM. The catch trial responses were not used to update the Psi staircases, yet ensured that once the staircase had converged, subjects still occasionally received faster or slower ground-truth trials. In Session 2, for each modality (Interoception, Exteroception) we used 12 catch trials, along with 48 Psi trials, for a total of 60 trials. In general, these changes improved the stability and reliability of staircase convergence (see Supplementary **Fig. 2.e**).

### Analysis

#### Statistical Analysis and Software

For both Session 1 and 2, we conducted planned statistical comparisons of threshold, slope, confidence ratings, meta-d and SDT parameters (*d*’ & M-ratio), across the two modalities using paired-samples t-tests at each timepoint separately. We further conducted an apriori assessment of test-retest reliability using the Pearson correlation coefficient for threshold and slope between Session 1 and 2. We further hypothesized that HRD thresholds would correlate with heartbeat counting scores, assessed via a priori correlation analysis. In addition to these planned analyses, we conducted exploratory group by time repeated measures ANOVAs on HRD parameters to assess possible interaction effects, and also estimated exploratory cross-correlation matrices for all HRD and HBC parameters at both time points.

Statistical analyses were conducted using Pingouin v0.3.9 (Vallat, 2018). The Bayes Factors were computed using a Cauchy scale of 0.707 and the p-values for the 2-way repeated measure ANOVAs were adjusted using the Greenhouse-Geisser correction. Correlation coefficients were tested using skipped correlations as implemented in Pingouin, which are robust to outliers (Pernet et al., 2013). We controlled for multiple comparisons in the correlation matrices using FDR correction (p_FDR_ < 0.01). Where applicable, outliers were detected and rejected using the absolute deviation around the median rule (Leys et al., 2013). Test-retest reliability was tested using the Pearson correlation from the same package. Figures were created using Matplotlib (Hunter, 2007), Seaborn (Waskom et al., 2020) and Arviz (Kumar et al., 2019). Distributions for repeated measures are represented using an adaptation of raincloud plots (Allen et al., 2021). All preprocessed data and analysis scripts supporting the results described in this paper are available at https://github.com/embodied-computation-group/CardioceptionPaper.

#### Bayesian Modelling of Psychometric Functions

Although Psi adaptively estimates slope and threshold parameters at each trial, we elected to apply a post hoc modelling approach to improve psychometric estimation. The post hoc modelling was applied after rejecting trials with extremely fast (< 100ms) responses or during epochs containing unreliable cardiac signals. In general, this approach yielded highly similar results as the Psi estimates - see supplementary materials (**Supp Fig. 1 & 2**) for a comprehensive analysis. We used the absolute deviation around the median rule (Leys et al., 2013) to identify and reject outliers in the instantaneous heart rate time series. Trials were rejected if the average of the heart rate was considered an outlier, or if the standard deviation of the pulse-to-pulse intervals was detected as outliers when compared to the other trials. This ensured that we only included responses in which the participant was in principle able to correctly estimate their cardiac frequency. We implemented Bayesian modelling of the psychometric functions using PyMC3 (Salvatier et al., 2016, p. 3). We used the NUTS sampling algorithm (Hoffman & Gelman, 2011) to update and estimate the posterior probability of the slope (β) and threshold (α) parameters (n_chains_=2, n_tuning_=4000, n_samples_=1000) for each subject and modality separately. We used a cumulative normal function so the results can be compared to what is estimated by the Psi staircase (see Supplementary Material **Fig. 2**). The psychometric model parameters were defined as:

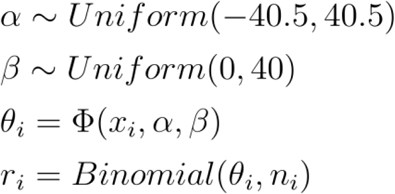

Considering the *i*th intensity levels, for the trials with a stimulus intensity *x_i_* we observed a total of *n_i_* responses, among which *r_i_* were “Faster“ responses. Here, **ϕ** is the cumulative normal function defined by:

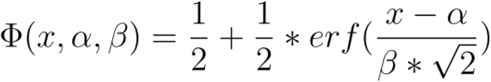

Because we aimed to correlate the resulting scores with other variables and were not interested in group-level means, we fitted the model for each subject and each modality separately in a non-hierarchical manner. These post hoc models are included in the cardioception toolkit, and future releases will provide easy to use hierarchical group estimation, to facilitate for example between groups analyses (Valton et al., 2020). All subsequent psychometric behavioural analyses were performed on the post hoc estimated parameters.

#### Signal Theoretic Modelling of Perceptual and Metacognitive Sensitivity

For these analyses, accuracy was coded such that a “Faster” response was correct only when the intensity **Δ**-BPM was greater than 0, and a “Slower” response was correct only when the intensity **Δ**-BPM was smaller than 0. Confidence ratings were binned into 4 equally spaced bins before modelling using the discreteRatings() functions from metadPy, a custom python package for metacognition modelling (https://github.com/LegrandNico/metadPy). We used a standard Signal Detection Theory (SDT) approach to estimate type 1 (i.e., perceptual) and type 2 (i.e., metacognitive) bias and sensitivity from the binned confidence ratings. Briefly, this model operationalizes metacognitive “insight” as the sensitivity of subjective confidence ratings to ground truth accuracy; e.g., by defining a receiver-operating characteristic (ROC) curve relating metacognitive “hits” - *p*(*Confidence* = *High*|*Response* = *Correct*) - and “misses” - *p*(*Confidence* = *High*|*Response* = *Incorrect*) - (Fleming & Lau, 2014; Maniscalco & Lau, 2012a). This measure is known as meta-*d*’, and is an index of metacognitive sensitivity akin to *d*’. However, as meta-d is known to be influenced by overall *d*’, and interoceptive accuracy is generally lower than exteroceptive on our task, we analyzed the parameter M-ratio (meta-*d*’/*d*’), also known as “metacognitive efficiency”. This parameter operationalizes metacognitive insight in signal theoretic units; e.g., the proportion of available sensory evidence utilized by the subjective confidence response. Perceptual and Metacognitive parameters were estimated using an adapted hierarchical Bayesian model from the HMeta-d toolbox (Fleming, 2017). We reparameterized this model to implement a paired-samples t-test estimating the within-subject impact of modality (interoceptive vs. exteroceptive) on M-ratio. The significance of this effect was then assessed by checking if the 94% highest density interval (HDI_94%_) includes zero or not.

#### Physiological Analysis

The time-series recorded through photoplethysmography (PPG) were analysed using Systole v0.1.3 (Legrand & Allen, 2021). The PPG signal, sampled at 75 Hz, is a measure of peripheral blood oxygenation level, in which cardiac cycles can be tracked by detecting abrupt increases following cardiac contraction and blood circulation (i.e., systolic peaks). The signal was first resampled to 1000Hz using linear interpolation. This procedure simplifies the measurement of the pulse-to-pulse intervals and can refine the peak detection precision, and the resulting heart rate when the initial sampling rate is low (Quintana et al., 2016). Clipping artefacts were removed using cubic spline interpolation (van Gent et al., 2019), the signal was then squared for peak enhancement and normalized using the mean + standard deviation using a rolling window (window size: 0.75 seconds). All positive peaks were labelled as systolic (minimum distance: 0.2 seconds). This procedure was applied both for the online heart rate recording during the Heart Rate Discrimination task (segments of 5 seconds) and for the Heartbeat Counting task. If any interbeat interval higher than 120 BPM or lower than 40 BPM was detected during the online recording of the Heart Rate Discrimination task, an error message was presented on the screen to ask the participant to stay still, and the trial was started again, up to 5 times consecutively before dropping the trial. As the correct detection of heartbeats is critical for the Heartbeat Counting task, we ran additional artefacts correction steps to control for erroneous or missed detection of some heartbeats. Extra heartbeats (i.e., erroneous labelling of peaks in PPG signal) were automatically removed using an artefact correction algorithm (Lipponen & Tarvainen, 2019) implemented in Systole (Legrand & Allen, 2021). All raw time series were manually inspected to ensure correct systolic peak detection. The HTML reports detailing these preprocessing steps of the HRD and the HBC tasks are made available online with the GitHub Repository associated with this paper.

#### Heartbeat Counting Analysis

We derived an accuracy score following previous recommendations (Garfinkel et al., 2015; Hart et al., 2013) as follows:

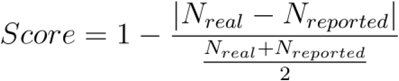

This score has a maximum of 1 and indicates the similarity between the objective recorded number of heartbeats and the number reported by the participant (a score of 1 indicating a perfect match). We used the absolute deviation around the median rule (Leys et al., 2013) to automatically detect and remove extreme responses that are more likely to reflect erroneous numbers provided by the participant. The remaining scores were subsequently averaged for each participant.

## Results

### Characterizing the Interoceptive and Exteroceptive Psychometric Function

We first analyzed the threshold (**α**) and slope (**β**) psychometric parameters using the estimates from the post hoc Bayesian model, separately for both sessions (see **Methods** for more details and **Supplementary Results** for comparison of Psi and post hoc parameters). These analyses serve to both characterize the overall shape of the two psychometric functions, which can inform the setting of prior parameters in future experiments, and assess how belief accuracy, bias, and precision differed between the two conditions. Further, to explore possible Session by Modality interactions, we fit repeated measures ANOVAs to each parameter of interest.

A paired sample *t-test* at Session 1 revealed that threshold was significantly lower in the interoceptive condition than in the exteroceptive condition (mean_Intero_ = −6.97, CI_95%_ [−8.59, −5.37], mean_Extero_ = 1.36, CI_95%_ [0.9, 1.85], *t*_(205)_ = −9.89, *p* < 0.001, BF_10_ = 1.15e+16, *d* = −0.93), indicating a robust negative bias; i.e., heart rate underestimation. This effect was replicated at Session 2 (mean_Intero_ = −8.50, 95% CI_95%_ [−10.09, −6.91], mean_Extero_ = 0.008, 95% CI [−0.48, 0.50], *t*_(190)_ = −11.15, *p* < 0.001, BF_10_ =2.85e+19, *d* = −1.03). We further noted a marked increase in inter-subject variance for interoceptive thresholds, range_Intero_ = [−38.3, 34.0] vs. exteroceptive thresholds, range_Extero_ = [−9.8, 16.39], indicating substantially more inter-individual variance in the magnitude of interoceptive biases. In contrast, when compared to a null hypothesis of 0 bias, follow-up one-sample *t-tests* on exteroceptive thresholds revealed a slight but highly significant positive bias at Session 1 (mean = 1.39, CI_95_ = [0.9, 1.89], *t*_(203)_=5.58, *p* < 0.001, BF_10_ = 1.28e+05, *d* = 0.99), which was not present at Session 2, in which we observed instead a strong evidence for an absence of difference (mean =0.01, CI_95_ =[−0.48, 0.52], *t*_(189)_ = 0.06, *p* = 0.94, BF_10_ = 0.08, *d* = 0.05). Finally, exploratory repeated measures ANOVA revealed significant main effects of Session (F_(1,178)_ = 13.20, *η*_p_2 = 0.06, *p* < 0.001) and Modality (F_(1,178)_ = 127.53, *η*_p_2 = 0.41, *p* < 0.001), indicating that thresholds were significantly reduced across sessions for both modalities, and that interoception was more biased across both sessions, but with no Session by Modality interaction (F_(1,178)_ = 0.60, *η*_p_2 = 0.003, *p* = 0.43). See **Fig. 2** and **6B** for illustration of these effects. Collectively these results show that interoceptive heart rate beliefs are robustly biased towards underestimation, and show greater inter-individual variance, than the exteroceptive control condition.

**Legend Figure 2:**
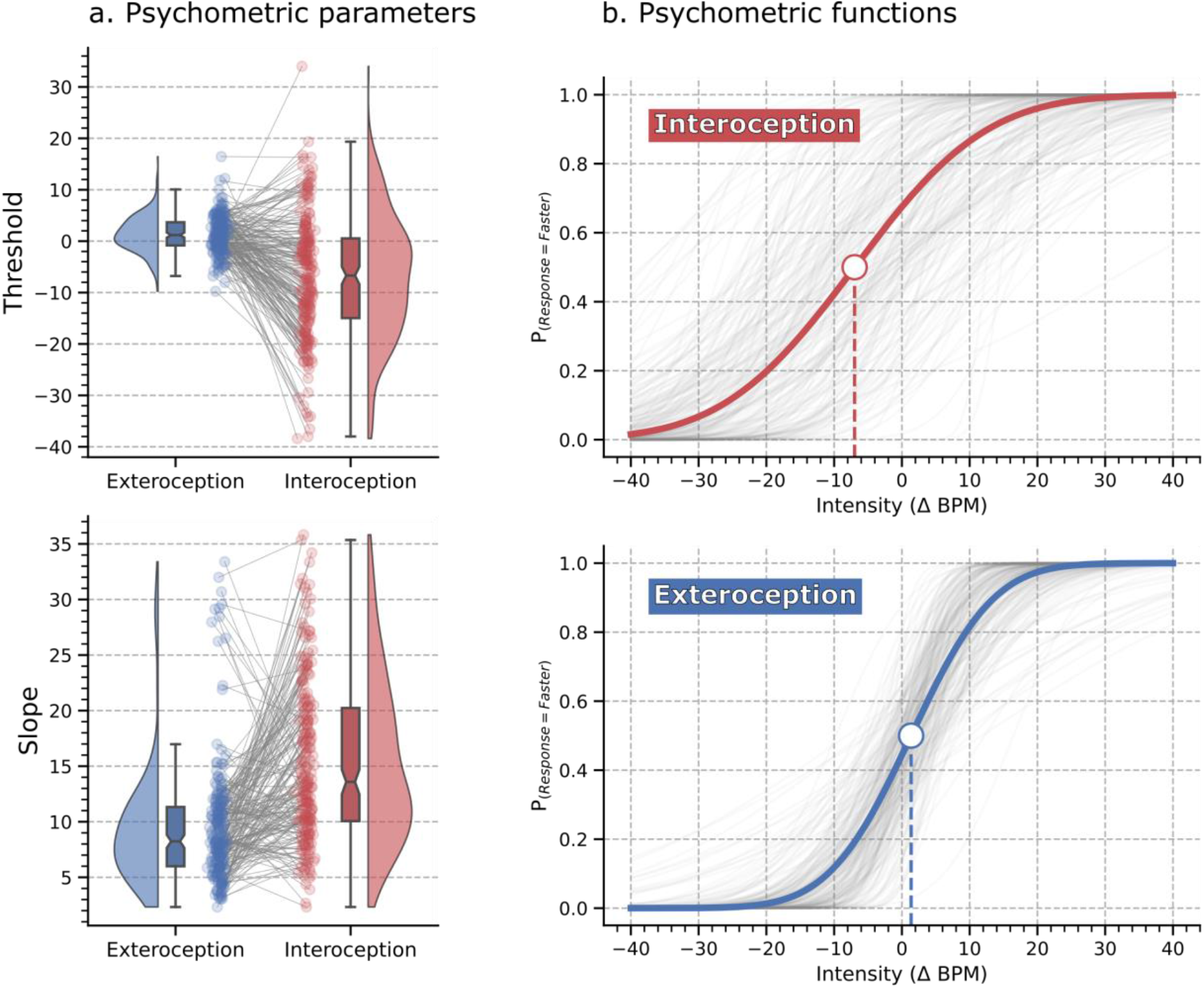
Psychometric parameter estimates and fitted interoception and exteroception psychometric functions (Session 1). **A.** Repeated measures raincloud plots visualizing threshold and slope parameters of the psychometric functions across the two modalities (interoception and exteroception). Data points for every individual are connected by a grey line to highlight the repeated measure effect. **B.** The grey lines show individual subject fits. The dark red and blue lines show the grand mean psychometric function, depicting averaged threshold and slope. Grand mean thresholds are marked by the large point, where the psychometric function crosses 0.5 on the ordinate axis. We observed a strong effect of interoception on both slope and threshold as compared to the exteroceptive control condition. The negative bias observed on threshold demonstrates that participants underestimated their heart rate on average. The greater slope indicates a less precise decision process.

We next consider the slope of the interoceptive and exteroceptive functions. While the threshold indicates the overall accuracy and bias of the decision-making process, the slope characterizes the precision or uncertainty of this process. A higher slope indicates a less steep (i.e., more shallow) psychometric function, indicating lower precision (higher uncertainty) for that condition. A paired sample *t-test* revealed that slope was significantly higher in the interoceptive condition as compared to the exteroceptive (mean_Intero_ = 15.34, CI_95%_ [14.39, 16.36], mean_Extero_ = 9.58, CI_95%_ [8.89, 10.39], *t*_(205)_ = 9.05, *p* < 0.001, BF_10_ = 4.97e+13, *d* = − 0.88). This effect was reproduced in Session 2, with interoceptive slope again greater than exteroceptive slope at retest (mean_Intero_ = 11.96, 95% CI [11.17, 12.76], mean_Extero_ = 8.69, 95% CI [8.11, 9.26], *t*_(190)_ = 7.29, *p* < 0.001, BF_10_ = 9.12e+08, *d* = 0.67). Exploratory repeated measures ANOVA further revealed main effects of Session (F_(1,178)_ = 31.27, *η*_p_2 = 0.14, *p* < 0.001) and Modality (F_(1,178)_ = 106.29, *η*_p_2 = 0.37, *p* < 0.001), as well as an interaction between these two factors (F_(1,178)_ = 9.46, *η*_p_2 = 0.05, *p* = 0.002). Thus, interoceptive slope showed a greater reduction across sessions (*t*_(178)_ = −5.35, *p* < 0.001, BF_10_ = 4.13+04, *d* = −0.53) than exteroceptive slope (*t*_(178)_ = −2.19, *p* = 0.029, BF_10_ = 0.875, *d* = −0.20). Collectively, these results demonstrate that interoceptive beliefs are less precise than exteroceptive. Further, we could hypothesize that interoceptive precision is more sensitive to practice and training effects than exteroceptive precision. This notion cannot be fully tested here due to the methodological differences that we introduced between the two sessions. See, however, **Fig. 2** and **Supplementary Fig. 1** for the comparison between the two conditions.

### Perceptual and Metacognitive Sensitivity

In addition to the psychometric function underlying the subjective decision process, we also compared objective overall perceptual and metacognitive sensitivity between interoception and exteroception. To assess between condition differences on these indices, we performed paired-sample t-tests comparing interoceptive and exteroceptive performance on each key type 1 and type 2 measure (*d*’, average confidence, and M-ratio), as well as exploratory Modality by Session repeated measures ANOVAs on these variables.

The *d*’, which reflects discrimination sensitivity, was signifigantly lower in the interoception condition as compared to the exteroception condition during Session 1 (mean_Intero_ = 1.43, CI_95%_ = [1.34, 1.52], mean_Extero_ = 2.05, CI_95%_ = [1.96, 2.14]), *t*_(203)_ = −9.42, *p* < 0.001, BF_10_ = 5.06e+17, *d* = −0.97). We replicated this effect in Session 2 (mean_Intero_ = 1.87, CI_95%_ = [1.79, 1.96], mean_Extero_ = 2.25, CI_95%_ = [2.21, 2.3], *t*_(185)_ = −7.98, *p* < 0.001, BF_10_ = 4.45e+10, *d* = −0.77). We also performed an exploratory Session by Modality repeated measures ANOVA on these measures to assess overall interactions between these factors. We observed significant effects of both Session (F_(1,176)_ = 59.82, *η*_p_2 = 0.25, *p* < 0.001), Modality (F_(1,176)_ = 114.81, *η*_p_2 = 0.39, *p* < 0.001), and a Session by Modality interaction (F_(1,176)_ = 7.19, *η*_p_2 = 0.03, *p* = 0.008). This result shows that across both conditions, sensitivity increased from Session 1 to 2, with the greatest increase being observed in the interoceptive condition.

We next analyzed average subjective confidence, an indicator of metacognitive bias. We found that confidence ratings were significantly lower during the interoception condition as compared to the exteroception condition, both during the first session (mean_Intero_ = 51.52, CI_95%_ = [49.16, 53.87], mean_Extero_ = 61.44, CI_95%_ = [59.51, 63.57], *t*_(203)_ = −10.01, *p* < 0.001, BF_10_ = 2.3e+16, *d* = −0.62) and the second session (mean_Intero_ = 57.47, CI_95%_ = [55.41, 59.77], mean_Extero_ = 64.27, CI_95%_ = [62.4, 66.03], *t*_(189)_ = −7.15, *p* < 0.001, BF_10_ = 4.18e+08, *d* = −0.49). An exploratory repeated measures ANOVA revealed a main effect of Session (F_(1,176)_ = 26.37, *η*_p_2 = 0.13, *p* < 0.001), Modality (F_(1,176)_ = 101.37, *η*_p_2 = 0.36, *p* < 0.001) and a Session by Modality interaction (F_(1,176)_ = 8.72, *η*_p_2 = 0.04, *p* < 0.003). The average confidence was higher in the second session as compared to the first one (*t*_(176)_ = 5.13, *p* < 0.001, BF_10_ = 1.52+04, *d* = 0.30), and this increase was larger for the interoceptive condition (*t*_(176)_ = 6.02, *p* < 0.001, BF_10_ = 9.68+05, *d* = 0.36) than for the exteroceptive condition (*t*_(176)_ = 2.54, *p* < 0.01, BF_10_ = 1.93, *d* = 0.18). Overall, confidence was generally lower for interoceptive vs. exteroceptive confidence.

To assess metacognitive sensitivity for both modalities, we estimated metacognitive efficiency using hierarchical modelling of M-ratio (meta-*d*’/*d*’) (Fleming & Lau, 2014; Maniscalco & Lau, 2012a). We observed that the individual estimated M-ratio values, as estimated by the repeated measure model, were lower during the interoception condition (mean_Intero_ = 0.81, CI_95%_ = [0.78, 0.86]) as compared to the exteroception condition (mean_Extero_ = 0.96, CI_95%_ = [0.92, 1.01], see **Fig. 3.b**). This tendency is confirmed by inspecting the posterior distribution of the log-transformed repeated measure effect (mean = −0.19, HDI_94%_ = [−0.36, −0.06], see **Fig. 3.c**). Because the M-ratio reflects the relation between the amount of evidence for metacognitive judgement and the amount of evidence for the objective decision, our results suggest that 19% of the interoceptive evidence used for decision in the type 1 task is lost during the metacognitive evaluation of confidence, compared to just 4% evidence loss for exteroception.

**Legend Figure 3:**
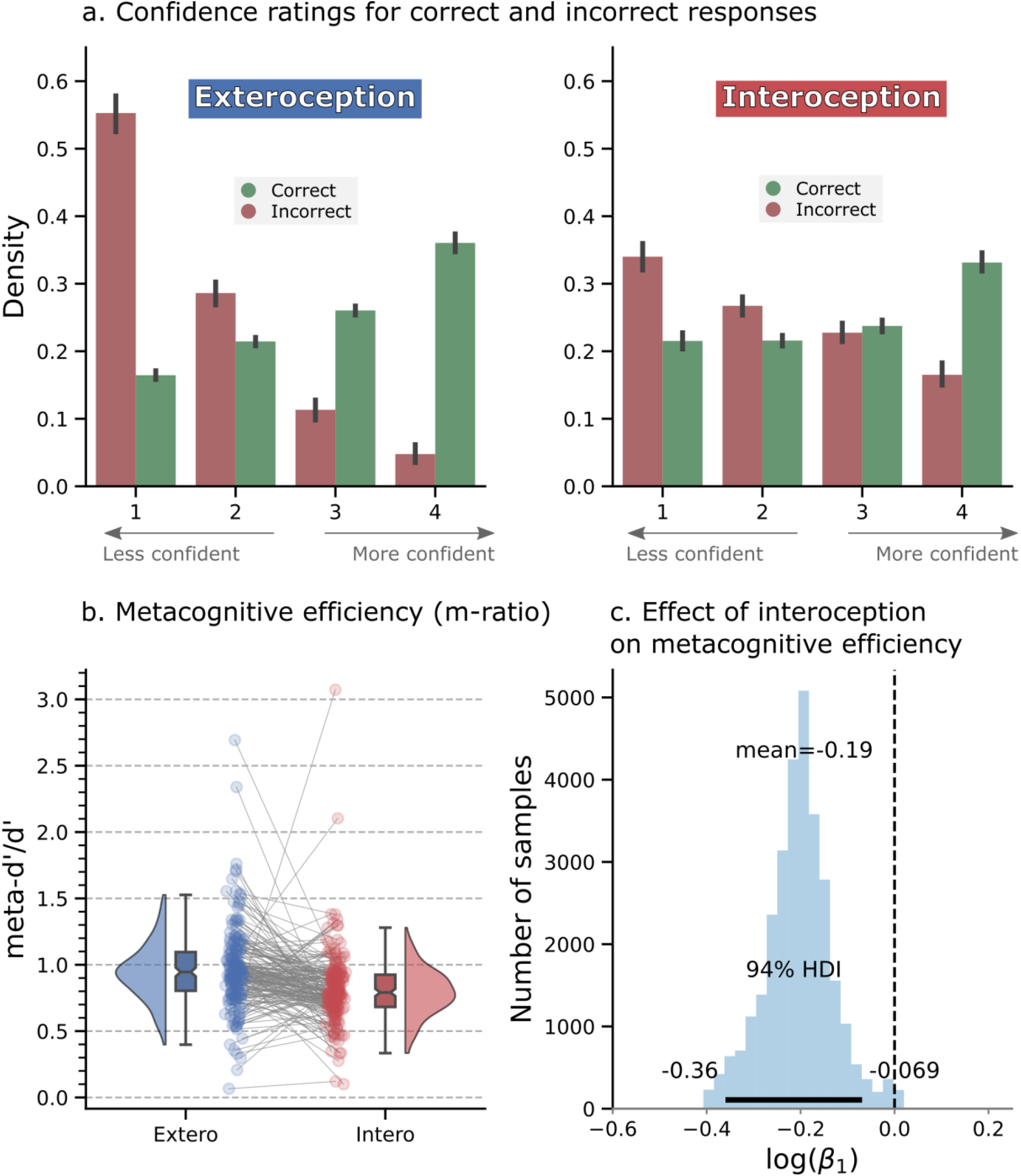
Visualization of metacognitive performance for interoception and exteroception conditions (Session 1). **A.** Histogram showing the distribution of binned confidence ratings for correct (green) vs. error (red) trials. Higher bins represent higher confidence ratings. Overall, participants were significantly less confident in the interoceptive condition and showed reduced metacognitive sensitivity as indicated by the flattening of the confidence distributions. **B.** To quantify this effect, we estimated “metacognitive efficiency”, a signal theoretic model of introspective accuracy which controls for differences in type 1 (discrimination) performance. Here, an M-ratio of 1 indicates optimal metacognition according to an ideal observer model, whereas values lower than this indicate inefficient use of the available perceptual signal. This model demonstrated that metacognitive efficiency was substantially decreased for interoceptive relative to exteroceptive judgements. **C.** Histogram of posterior samples from the beta value encoding the difference of interoception-exteroception in the repeated measures hierarchical model.

We replicated this finding in Session 2, where interoception M-ratio estimates were again lower (mean_Intero_ = 0.83, CI_95%_ = [0.8, 0.87]) than those for exteroception (mean_Extero_ = 0.96, CI_95%_ = [0.92, 1.01]), as well as in the posterior distribution of the repeated measure effect (mean = −0.17 HDI_94%_ = [−0.26, −0.03], see **Supplementary Material, Fig. 3**).

### Cross-modal Correlations

To investigate the construct validity of HRD performance measures, we conducted an exploratory correlation analysis relating individual differences in perceptual and metacognitive performance within and between the interoceptive and exteroceptive modalities. For this analysis, we refitted the meta-*d*’ model (Fleming, 2017) separately to each participant (i.e., in a non-hierarchical model), and extracted individual M-ratio values. Here, we sought to verify whether threshold, slope, or other type 1 or type 2 parameters were correlated across the two conditions. For example, if HRD performance primarily indexed general temporal estimation ability, we would expect a high correlation between interoceptive and exteroceptive thresholds, as well as with other type 1 performance variables. Alternatively, if participants used additional information, such as afferent cardiac sensory information and/or prior beliefs specifically about the heart rate, then we would expect little to no correlation between these parameters. Additionally, previous studies found that interoceptive metacognition is typically uncorrelated to exteroceptive metacognition, suggesting unique inputs for these self-estimates (Garfinkel et al., 2016). However, more recent work suggested the existence of a “metacognitive g-factor” indexed by high inter-modal correlations in metacognitive ability (Mazancieux et al., 2020; Rouault et al., 2018). We, therefore, included both type 1 measures (i.e., threshold, slope, *d*’, response time, and criterion) and type 2 measures (confidence, meta-*d*’, M-ratio) in one exploratory between-subject correlation analysis to probe the degree of within and between modality overlap in parameter estimates. To do so, we performed robust pairwise correlation tests between exteroception and interoception task parameters (Pernet et al., 2013), using a skipped correlation approach and correcting for multiple comparisons using a false-discovery rate (FDR, p_FDR_ < 0.01) correction. The resulting Spearman’s r coefficients for Session 1 are summarized in **Fig. 4.**

**Legend Figure 4:**
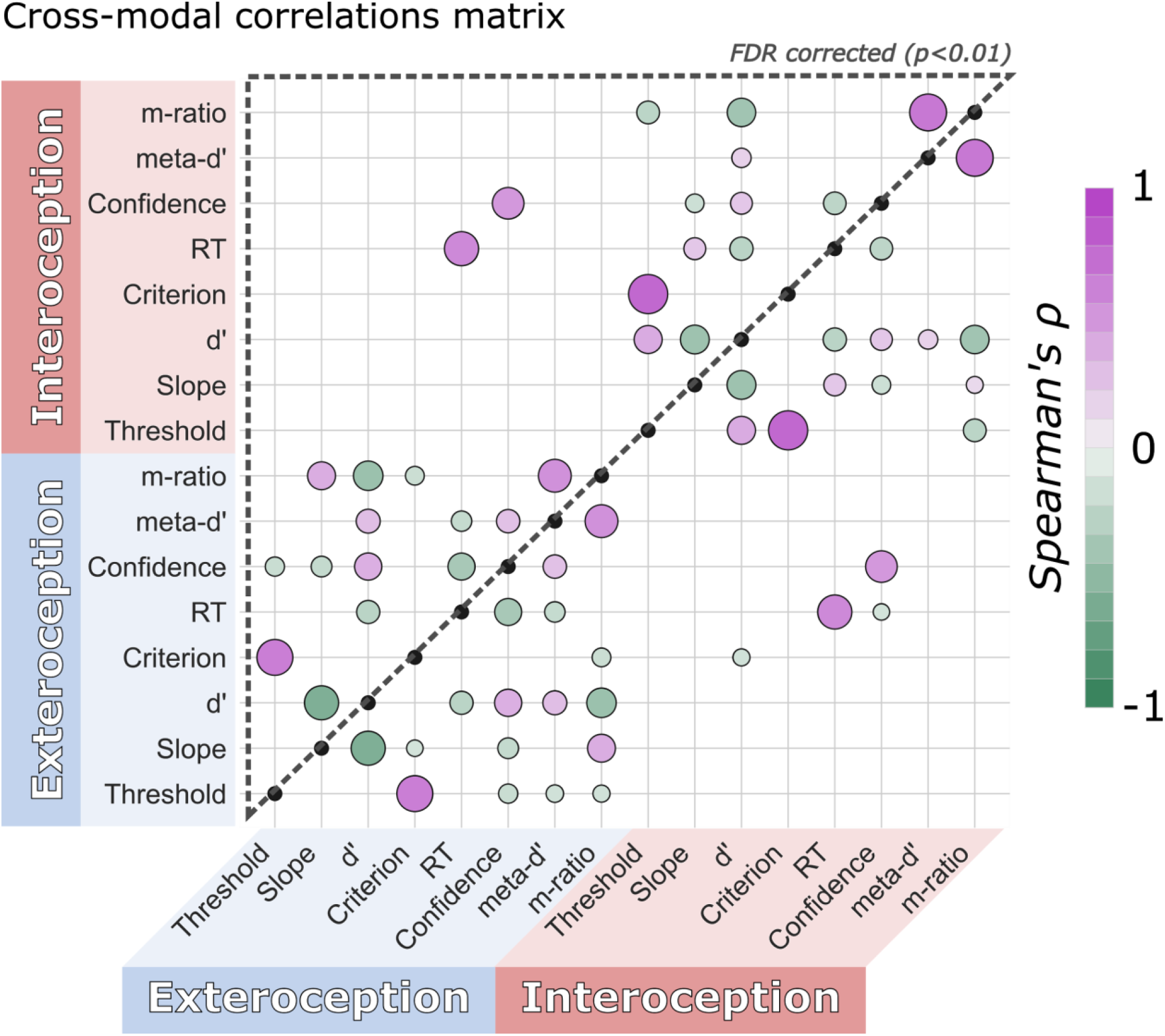
Cross-modal correlation heatmap of task parameters for interoception and exteroception conditions (Session 1). Overall, we observed that behavioural results were correlated within modalities but with limited dependence across modalities, the only exceptions were confidence and response time (RT). Only significant skipped Spearman correlations are represented. **The upper triangle only shows results surviving FDR correction (p_FDR_ < 0.01), while the lower right triangle of the matrix shows the uncorrected comparisons**. Colour and size of individual points indicate the sign and strength of estimated correlation coefficients. See supplementary Fig. 4 for Session 2 cross-correlations.

We observed more robust and consistent correlations between task parameters within each modality (interoception or exteroception), but few significant correlations between task modalities, indicating a high degree of independence between performance on the two task conditions. Interestingly, with the exception of reaction time, type 1 performance was largely uncorrelated between modalities, whereas at the metacognitive level only subjective confidence was highly correlated (r_s_ = 0.60, CI_95%_ = [0.52, 0.69], p < 0.001, n = 204, n_outliers_ = 5). These results may suggest that individuals use similar “self-beliefs” about their performance on both task modalities (Fleming & Daw, 2017). A similar overall pattern was observed in Session 2, albeit with a modest but significant relationship between interoceptive and exteroceptive thresholds (r_s_ = 0.26, CI_95%_ = [0.13, 0.39], *p* < 0.001, n = 190, n_outliers_ = 6, see **Supplementary Results** for the full correlation matrix).

### Correlation with the heartbeat counting task parameters

As a final check of construct validity, we assessed how our new task relates to the standard heartbeat counting task. We thus correlated HRD performance variables (psychometric thresholds and slopes) with the HBC scores. We found that the interoceptive thresholds from Session 1 were positively correlated with the global heartbeat counting score (see **Fig. 5.a**; r_s_ = 0.29, CI_95%_ = [0.16, 0.42], *p* < 0.001, n = 193, n_outliers_ = 1). No significant correlation was found between the heartbeat counting score and the exteroceptive threshold (r_s_ = −0.04, CI_95%_ = [−0.18, 0.1], *p* = 0.58, n = 193, n_outliers_ = 13). We further replicated the correlation between HRD threshold and HBC iACC scores in Session 2 (r_s_ = 0.19, CI_95%_ = [0.05, 0.33], *p* = 0.01, n = 178, n_outliers_ = 0).

**Legend Figure 5:**
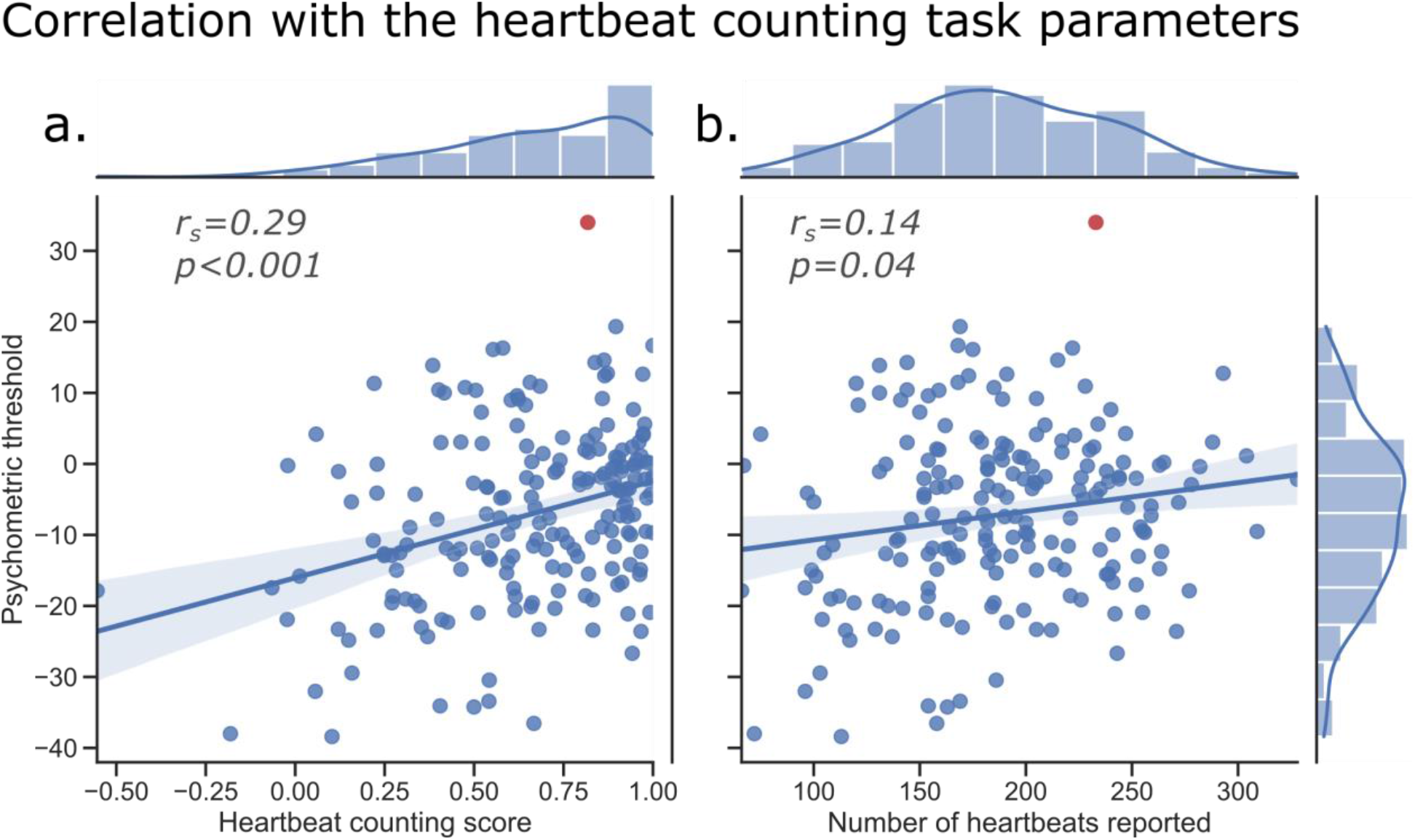
Correlation between the psychometric threshold and heartbeat counting performance. (Session 1) **A.** We found that heart rate discrimination (HRD) thresholds correlate positively with heartbeat counting (HBC) interoceptive accuracy scores. A lower threshold (i.e., a more negative bias) on the HRD task was associated with lower performance on heartbeat counting. We suggest that low scores on the heartbeat counting task are associated with a tendency to undercount the number of heartbeats. **B.** The psychometric threshold was associated with the total number of heartbeats reported during the heartbeat counting task. The correlation was also found while controlling for the heart rate during the task (not shown). These results suggest that participants’ inability to reliably count their heartbeats is partially explained by lower interoceptive thresholds. Outliers detected by the skipped correlation are reported in red. The r_s_ and p values are from the bootstrapped Spearman coefficient. The regression line is only fitted to non-outlier data points. The shaded area represents the bootstrapped confidence interval (95%).

The previous results suggest that the bias observed in the heartbeat counting task might be at least partially explained by the participants’ tendency to underestimate their own heart rate. To corroborate this notion, we attempted to verify the association between the psychometric threshold obtained during the HRD task, which quantifies the heart rate underestimation, and the total number of heartbeats reported by the participant during the HBC task (see **Fig. 5.b**). The psychometric threshold was positively correlated with the total number of heartbeats counted by the participants (r_s_ = 0.14, CI_95%_ = [0.01, 0.28], *p* = 0.04, n = 193, n_outliers_ = 1). It could be argued here that the actual heart rate of the participant may directly influence the total number of counted heartbeats, as the number of heartbeats that can be potentially counted naturally increases with increments in heart rate frequency. To control for this possible confound, we performed a semi-partial correlation between the psychometric threshold and the total number of counted heartbeats while controlling for the relation between the number of counted heartbeats and the number of actual heartbeats detected in the PPG signal. This analysis revealed a positive correlation between these two variables (r_s_ = 0.20, CI_95%_ = [0.07, 0.34], *p* = 0.004, n = 193, n_outliers_ = 2).

### Reliability of psychometric parameters

Intrinsic or test-retest reliability is a critical feature of any measurement, in particular, if it is to be useful for clinical diagnostic or intervention purposes. To evaluate reliability, we calculated the correlation coefficient for Session 1 and 2 interoceptive thresholds and slopes, obtained on average 46.79 days apart from each other. Threshold was highly correlated between sessions, showing good reliability (r = 0.51, *p* < 0.001, BF_10_ = 5.04e+10, see **Fig. 6**). In contrast, Slope was not correlated across sessions (r = 0.10, *p* = 0.15, BF_10_ = 0.25), potentially indicating a poor reliability of this parameter.

**Legend Figure 6:**
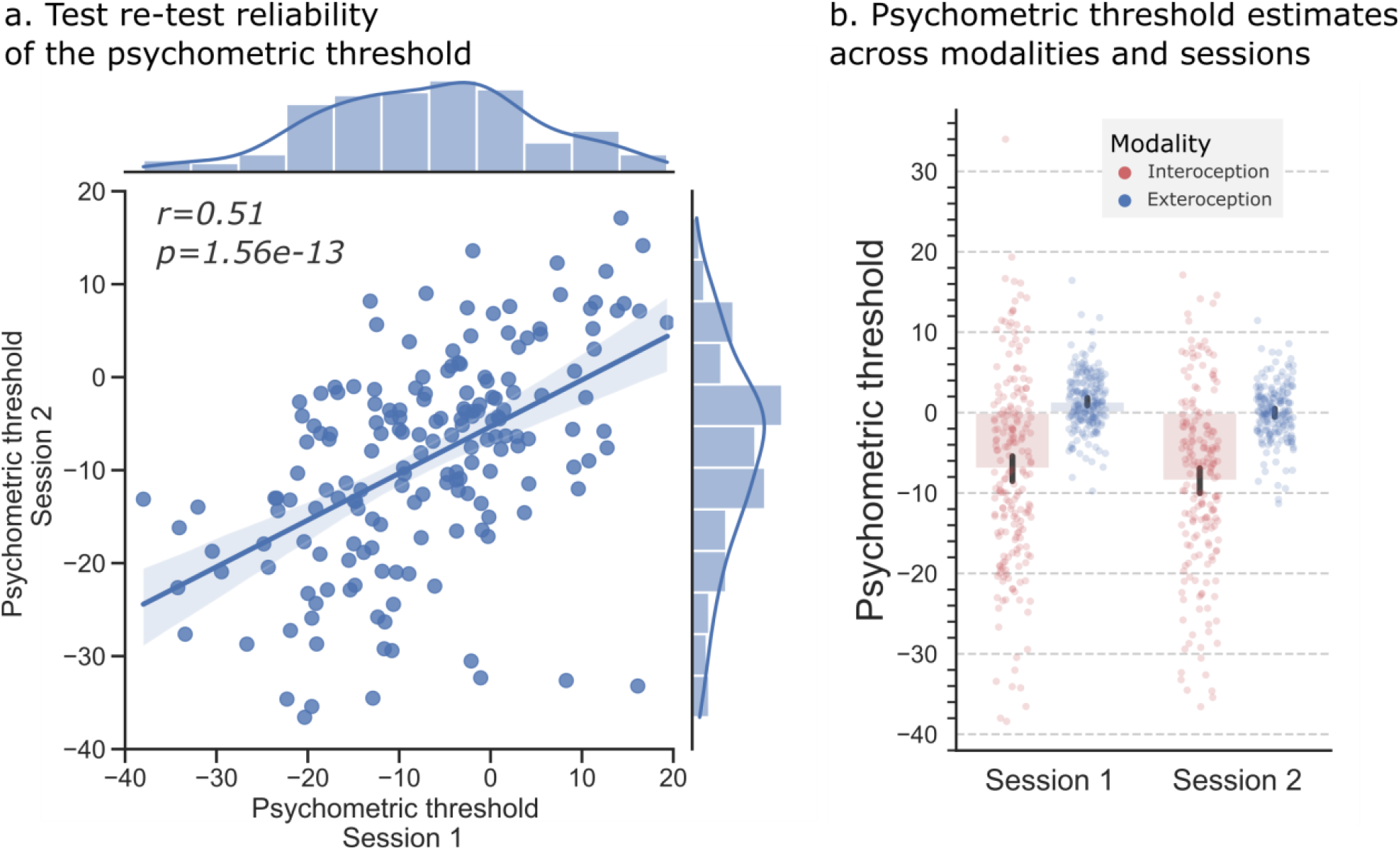
Test-retest reliability of the psychometric threshold. The psychometric threshold estimated using a Bayesian post hoc approach provided correct test-retest reliability. **A.** Correlation between the interoception threshold estimates in Sessions 1 and 2. Outliers detected by the skipped correlation were removed and the reliability was tested using a Pearson correlation. The r_s_ and p values were calculated using the bootstrapped Spearman coefficient. The regression line was only fitted to non-outlier data points. The shaded area represents the bootstrapped confidence interval (95%, 1000 iterations). **B.** Distribution of threshold Bayesian estimates across sessions and modalities (n=204 for Session 1; n=190 for Session 2). The error bars represent the bootstrapped confidence interval (95%, 1000 iterations).

## Discussion

The measurement of cardiac interoception is a methodological puzzle that has challenged generations of psychologists and psychophysiologists (Ainley et al., 2020; Brener Jasper & Ring Christopher, 2016; Chen et al., 2021; Zamariola et al., 2018; Zimprich et al., 2020). Here, we suggest that this difficulty arises in part from a reluctance to treat subjective perceptual beliefs about the heart rate as a core component of interoception. To remedy this gap, we introduce a new Heart Rate Discrimination (HRD) task, which incorporates a Bayesian psychophysical procedure for measuring the accuracy, precision, and metacognitive sensitivity of cardiac decisions. In a study of 223 healthy participants, we observed robust and consistent heart rate underestimation. We also found that interoceptive beliefs and metacognition are more imprecise as compared to the exteroceptive control condition. Our results indicate that interoceptive beliefs as measured by the HRD are not strongly correlated with other exteroceptive temporal beliefs, but share some variability with indexes of interoception measured by the Heartbeat Counting task. In general, these effects were robustly replicated across two testing sessions, with interoceptive thresholds, in particular, exhibiting good within-participant test-retest reliability. These features make the HRD well-suited for the measurement of interoceptive biomarkers in clinical populations, and for basic research probing the underlying mechanisms underlying cardiac beliefs and their influence on behaviour.

Our principal finding is that participants consistently underestimate their resting heart rate by 7 BPM on average, with substantial inter-individual variation around this value (**Δ**-BPM threshold range = [−39, 30]) (**Fig. 2**). This finding is consistent with repeated reports that heartbeat counting scores are driven by undercounting (Zamariola et al., 2018) − for discussion, see (Ainley et al., 2020; Corneille et al., 2020; Zimprich et al., 2020). We also find that interoceptive HRD thresholds are moderately correlated with HBC iACC scores, such that fewer counted heartbeats correlate with a lower HRD threshold (see **Fig. 5**). When comparing interoceptive and exteroceptive thresholds, we further found a similar positive correlation at session 2 (**see Supp Fig. 4**). These results highlight the unique sources of variance influencing interoceptive beliefs, such that HRD thresholds (and by extension, HBC scores) are likely to be driven by a combination of general temporal estimation ability, bottom-up cardio-sensory inputs, and top-down beliefs about the heart rate.

The ability to distinguish these contributions is a unique strength of the HRD. Future clinical investigations will benefit from including both interoceptive and exteroceptive conditions to tease apart these different potential causes of apparent interoceptive dysfunction. For example, if a participant group shows general cross-domain main effects on both interoceptive and exteroceptive thresholds, this would indicate a general deficit in temporal estimation rather than an alteration of interoceptive beliefs. In contrast, group or conditional interaction effects on the interoceptive threshold or slope, in the absence of any exteroceptive effects, would indicate a specific deficit in monitoring bodily sensations and updating cardiac beliefs. In this way, investigating conditions by group interactions on HRD parameters should hopefully improve the specificity of interoception research.

Another important finding is that interoceptive precision, as measured by the slope of the psychometric function, was substantially lower than exteroceptive precision (**Fig. 2 and Supp Fig. 1**). This is an interesting finding in light of recent theoretical and computational models which hypothesize that interoceptive sensory signals in the brain may generally be more imprecise than their exteroceptive counterparts (Ainley et al., 2016; Allen et al., 2019; Allen & Tsakiris, 2018). This hypothesis is based on influential “interoceptive predictive processing” models which emphasize the top-down, belief-driven nature of embodied self-perception. On these accounts, subjective interoceptive sensations are more likely to reflect the integration of top-down, prior expectations about the bodily self with ascending sensory inputs, with each signal weighted by their respective precision or confidence (Allen, 2020; Allen & Friston, 2018; Barrett & Simmons, 2015; Seth, 2013). The finding that interoceptive decisions are associated with lower precision may thus indicate that ascending cardiac signals are themselves inherently imprecise, or those prior beliefs encoding expected interoceptive precision are themselves more uncertain.

It should be noted however that “precision” as measured by the HRD indicates the uncertainty of the psychological decision process, and should not yet be treated as a direct read-out or measurement of the computational process by which prediction error signals are “precision-weighted”, which is thought to depend on neurobiological gain control (Bastos et al., 2012; Feldman & Friston, 2010). While previous investigations in the exteroceptive domain demonstrated a link between behavioural variability of this sort and neurocomputational precision (Eldar et al., 2013; Hénaff et al., 2020; van Bergen et al., 2015; Warren et al., 2016), in advance of direct evidence in the interoceptive domain this link should be interpreted with caution. Nevertheless, a unique benefit of our approach is that future studies could combine the HRD with computational modelling and direct neuronal recordings to conclusively establish the potential link between these parameters, and to tease apart the contributions of prior versus sensory precision to the imprecision observed here in heart-rate decisions (see e.g. Allen et al., 2019; Smith et al., 2020, 2021 for potential modelling applications).

Finally, we observed a robust reduction in metacognitive efficiency for interoceptive versus exteroceptive decisions. Although individual levels of subjective confidence (i.e., metacognitive bias) were highly correlated between modalities, metacognitive efficiency itself was not. This speaks to ongoing debates about the modularity of metacognition (Rouault et al., 2018), indicating that metacognitive ability in the interoceptive domain is largely unrelated to exteroceptive self-monitoring, in line with previous findings on this topic (Beck et al., 2019; Garfinkel et al., 2016). In light of these results, it is interesting to speculate as to the divergent mechanisms that might underlie metacognition in these two domains.

Numerous computational accounts emphasize that accurate metacognitive self-monitoring is likely to depend on a process by which the precision of the sensory signals underlying the type 1 decision is “read-out” by a higher-order metacognitive module, such that noisy, imprecise signals can be expected to degrade both perceptual performance and metacognitive sensitivity (Fleming et al., 2012; Maniscalco & Lau, 2016). However, other accounts emphasize that top-down “self-beliefs” may play a crucial role in shaping the interaction between low-level precision and higher-order metacognition (Allen et al., 2020; Fleming & Daw, 2017). Speculatively, our findings may suggest that in the cardiac domain, metacognition is largely dominated by top-down beliefs, rather than pure sensory read-out. Alternatively, if the reduced interoceptive precision observed here relates primarily to the uncertainty of cardiac sensory afferents, then this effect may be simply a result of the metacognitive system accurately reading out the low sensory precision. Teasing apart these different hypotheses through targeted causal manipulations of cardiac sensory signals and prior beliefs will hopefully shed new light on metacognitive insight into the bodily self.

### Strengths of the Heart Rate Discrimination Task

The HRD has several important methodological and practical strengths that support its utility in both basic and clinical research. First, the psychometric curve is estimated across trials relative to the ground truth heart rate. This allows us to differentiate the bias and precision of cardiac beliefs, in a way in which previous tasks such as HBC and HBD cannot. For example, it could be expected that the overall shape of the psychometric function may change under cardiovascular arousal, and the magnitude of this change could be an important marker of inter-individual differences in interoceptive reactivity.

A second feature of the HRD is the inclusion of an exteroceptive control condition, enabling measurements in the same units (**Δ**-BPM) in both modalities. This provision of sensible, easy to interpret units enables precise, meaningful comparisons across different studies, improving metric interpretability. The exteroceptive control condition itself has several additional benefits; it facilitates the use of the task in neuroimaging studies aiming to isolate more specific neural correlates of cardioceptive beliefs and allows for the differentiation of clinical symptoms into specific interoceptive deficits and more general temporal estimation deficits.

A third strength is that up to 100 HRD trials can be collected in as little as 25 minutes using standard physiological recording equipment. This is critical for clinical studies where testing time is often limited. A core contribution of the HRD is that it provides a novel decision axis through which researchers can probe interoceptive beliefs and percepts: the moment to moment decision of how fast one’s heart is beating. This trial design means that the HRD is amenable to a variety of quantitative modelling techniques such as hierarchical modelling of psychometric functions, or through computational modelling using reinforcement learning and similar approaches (Mathys et al., 2014; Petzschner et al., 2021). This feature facilitates testing mechanistic hypotheses about how cardioceptive beliefs are formed and updated and could be paired with, for example, the probabilistic manipulation of attention or performance feedback to delineate the role of prior beliefs and sensory prediction errors.

In general, we believe the HRD will be particularly useful as a clinical biomarker when comparing how specific populations update their cardiac beliefs under differing contexts − for example, one could test whether participants with anxiety show a tendency towards overestimating the heart rate at rest, or instead exhibit larger shifts in threshold and/or precision when comparing aroused vs. resting state performance.

### Limitations

The HRD offers several improvements to existing cardioceptive measures, including increased face validity, adaptability, and amenability to signal theoretic and other computational approaches to quantifying cardiac decisions. However, there are a few potential limitations of the task, and the results demonstrated here.

First, the HRD depends upon the online estimation of the heart rate within a five-second interval. While instantaneous measures of heart rate are generally robust, even within this time window there are likely to be within-trial shifts in high-frequency heart rate variability (HRV). Effectively this means that there is a theoretical lower bound on the precision with which one can estimate HRD thresholds, below which their interpretation becomes suspect. To control for this effect, we ensured that HRD step sizes (e.g., in terms of the minimum increment on **Δ**-BPM) are never lower than 1 BPM intervals, and also excluded trials with an extreme standard deviation of within-trial beat to beat intervals.

Another limitation is related to our implementation of the task as a two-interval forced-choice response. On each trial, participants first attended to their cardiac sensations and were then immediately presented with auditory feedback during the choice interval. This is a deliberate design decision, as the 2-IFC structure both ensures that participants have a window of interoception-only focus on each trial and renders the underlying behaviour more amenable to the signal theoretic assumptions of the metacognitive model (Galvin et al., 2003; Lee et al., 2018; Maniscalco & Lau, 2012b). We see this as an improvement over measures such as the heartbeat discrimination task, where subjects must perform a difficult simultaneous multisensory judgement, and it makes the task more amenable for identifying the neural or physiological correlates of HRD measures in the interoception-only time window. However, as a trade-off, this does induce a slight working-memory component to the task, as participants must form a belief about the heart rate and then hold it in mind while comparing it to the auditory feedback tones. This may be a limitation for studies comparing, for example, clinical populations with known working memory deficits. In this case, a variant of the task could easily be implemented in which the feedback tone is presented simultaneously with the listening interval, similar to recent tasks using a method of adjustment (Palmer et al., 2019).

The HRD task also includes an exteroceptive condition that has been designed to correspond as closely as possible to the interoceptive condition in terms of trial structure, timing and cognitive content that makes it appropriate for contrast-based analyses, e.g., in neuroimaging or physiological studies. It should be noted however that across trials, the frequency of the first “reference” stimulus is not derived from the heart rate but rather a random uniform distribution from 40 to 100 bpm. This means that the range of presented tones is greater in the exteroceptive vs interoceptive condition and that the exteroceptive psychometric function is essentially averaged across relatively slow and fast stimuli. If a participant has a large difference in responses across these bins, it could potentially limit the interpretation of the relative difference in interoceptive versus exteroceptive thresholds. One could alternatively generate these stimuli from a distribution matching that of the participants own heart rate, albeit with the trade-off of potentially feeding the participant implicit information about their heart rate. Future work should rigorously compare these possibilities to achieve optimal control over temporal and other cognitive confounds.

Finally, we do not present the HRD as measuring the objective sensitivity to ascending (i.e., baroreceptor mediated) cardiac sensations specifically. In the absence of further empirical data, interoceptive thresholds and/or precisions obtained by the HRD method should not be interpreted as a straightforward measure of the objective ability to discriminate viscerosensory sensations, as a variety of different strategies utilizing, for example, semantic beliefs or tactile inputs are likely to underlie decisions on the task, in particular under resting conditions (Khalsa et al., 2009). For researchers targeting specifically visceral ascending sensitivity, we would recommend approaches such as the MCS (Brener et al., 1993). Our task instead measures the bias and precision of subjective beliefs about the heart rate, which are likely to combine prior beliefs, contextual factors, and ascending (interoceptive and exteroceptive) sensory information where available. Future studies will pair causal manipulations of ascending cardiac signals with threshold measurement, to better delineate the degree to which these sensory inputs shape cardiac beliefs.

## Conclusion

In this study, we reported observations from the experimental use of the Heart Rate Discrimination task to measure the bias and precision of cardiac beliefs among a group of 223 individuals in a test-retest design. Our results have documented a robust tendency across participants to underestimate their heart rate, and have shown that interoceptive decisions are imprecise as compared to an exteroceptive control condition. We argue that the ability to objectively quantify these perceptual beliefs is a powerful tool for both basic and clinical interoception research. As this procedure is supported by psychophysics and Bayesian modelling of metacognition, it also calls for future methodological refinement and hypothesis-driven investigation to delineate the computational and physiological sources of cardiac beliefs.

## Acknowledgements

NL, NN, CMCC, MB, AS, NK, MN, and MA are supported by a Lundbeckfonden Fellowship (under Grant [R272-2017-4345]), and the AIAS-COFUND II fellowship programme that is supported by the Marie Skłodowska-Curie actions under the European Union’s Horizon 2020 (under Grant [754513]), and the Aarhus University Research Foundation. FF is supported by an European Research Council Starting Grant, under the European Union’s Horizon 2020 research and innovation programme (Grant agreement No. 948838). The authors further thank Benjamin Vincent for insightful discussions on the psychometric approach implemented here.

## Supplementary material

### Psychometric estimates using the Psi method

During Session 1, we used a 1-up/1-down procedure together with a Psi staircase to estimate the threshold of the psychometric function. During the first 30 trials of each condition, the intensity value was controlled by a 1-up/1-down staircase (Dixon & Mood, 1948) and the results were provided to the Psi staircase for initialization. We used this procedure to control for threshold convergence between the two techniques (results non reported here). During Session 2, we used only a Psi staircase procedure (Kontsevich & Tyler, 1999). The experimental setup was also slightly optimized between Sessions 1 and 2 (see **Material and Methods**). All these points could impact the efficiency and the parameters estimates of the Psi staircases. To check for possible deviation, we report in **Figure 1** of the **Supplementary Materials** the psychometric parameters estimates for slope and threshold across the two modalities and across the two sessions.

**Legend Supplementary Material 1:**
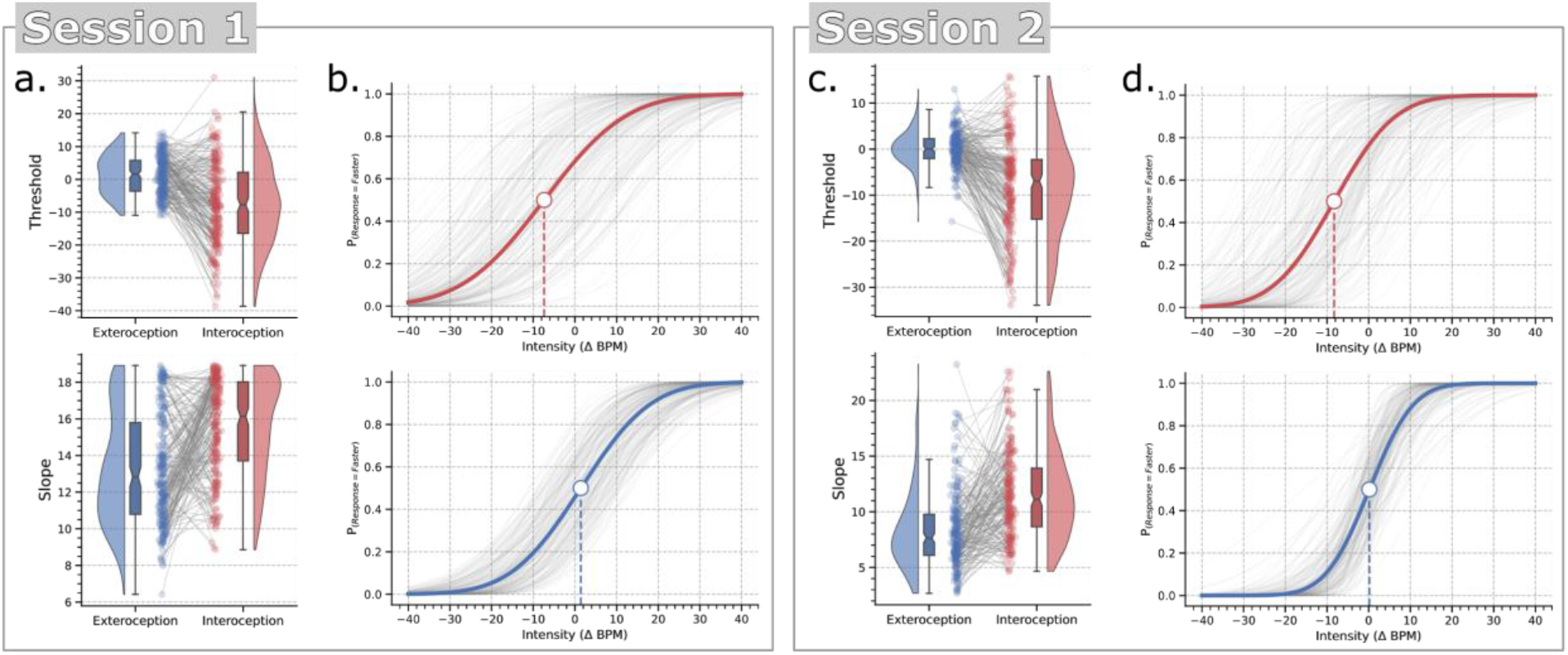
Psychometric parameters and psychometric functions estimated by the staircase using the Psi method from Sessions 1 and 2. Slope and threshold parameters of the psychometric functions for interoception (red) and exteroception (blue) conditions during Session 1 (n=206) (**A.**) and Session 2 (n=191) (**C.**). Psychometric functions fitted across interoceptive and exteroceptive conditions for Session 1 (**B.**) and Session 2 (**D.**). The grey lines show individual subject fits. The dark blue and red lines show the grand mean psychometric function, depicting the average threshold and slope. Both sessions show a strong effect of interoception on slope and threshold as compared to the exteroceptive control condition, with a negative bias and reduced precision for interoception.

Here, our results mirrored what we observed using the Bayesian estimates and comparing the two modalities conditions. We observed a bias in the interoceptive threshold as compared to the exteroceptive one in both Session 1 (t_(205)_ = −9.89, *p* < 0.001, BF_10_ = 1.20e+16, *d* = −0.90) and Session 2 (t_(190)_ = −11.66, *p* < 0.001, BF_10_ = 8.31e+20, *d* = −1.06). The slope, reflecting the imprecision of the decision, was also higher during interoception in both Session 1 (t_(205)_ = 7.86, *p* < 0.001, BF_10_ = 3.06e+10, *d* = 0.80) and Session 2 (t_(190)_ = 8.92, *p* < 0.001, BF_10_=1.50e+13, *d* = 0.86). Here, a higher slope reflects a less precise decision process. These results suggest that the two main psychometric effects (i.e., the threshold bias and slope increase during interoception) are robust and are not specific to one analytical approach in particular.

### Correlation between psychometric parameters estimated using the Psi method and a Bayesian post hoc model

In this paper, the psychometric parameters were estimated using a Bayesian model fitted on post-processed response data. This provides, in our opinion, a more robust framework for between session comparisons, and has the advantage to allow for behavioural and physiological data cleaning before model fitting. However, the values of the parameters can also differ between the Psi procedure and the final Bayesian estimates. We report in **Fig. 2** of the **Supplementary Materials** the relation between the values estimated by these two methods for both sessions.

When testing covariance using a Pearson correlation, we observed that the threshold estimates were highly consistent across the two estimation methods in both Session 1 (Exteroception: r = 0.63, CI_95%_ = [0.55, 0.71], n = 206; Interoception: r = 0.92, CI_95%_ = [0.91, 0.94], n = 206) and Session 2 (Exteroception: r = 0.96, CI_95%_ = [0.95, 0.97], n = 154; Interoception: r = 0.98, CI_95%_ = [0.98, 0.99], n = 147). These effects are illustrated in **Supplementary Material Fig. 2. a-c**.

We observed more variability in the estimation of slope, as reflected by the slightly lower correlation coefficients in Session 1 (Exteroception: r = 0.69, CI_95%_ = [0.62, 0.76], n = 206; Interoception: r = 0.80, CI_95%_ = [0.75, 0.84], n = 206) compared to Session 2 (Exteroception: r = 0.90, CI_95%_ = [0.87, 0.93], n = 154; Interoception: r = 0.78, CI_95%_ = [0.71, 0.84], n = 147). Notably, a ceiling effect and a systematic shift of the slope estimates was observed on Session 1 (see **Supplementary Material Fig. 2. b-d**). The ceiling effect was corrected in Session 2 by using larger parameter ranges. Here, the Bayesian approach included a larger prior range and was able to infer different slope values when the maximum was reached.

This analysis illustrates the power of a simple *post hoc* Bayesian modelling approach to improve and correct potential issues in the settings of the Psi staircase. This approach can be further expanded in future works, for example using fully hierarchical (i.e., mixed-effects) Bayesian modelling across participants and groups, improving the estimation of conditional differences in threshold or slope values. This could enhance statistical power by pooling and it further limits the influence of unlikely or outlier responses through group shrinkage effects on the parameter estimates.

**Legend Supplementary Material 2:**
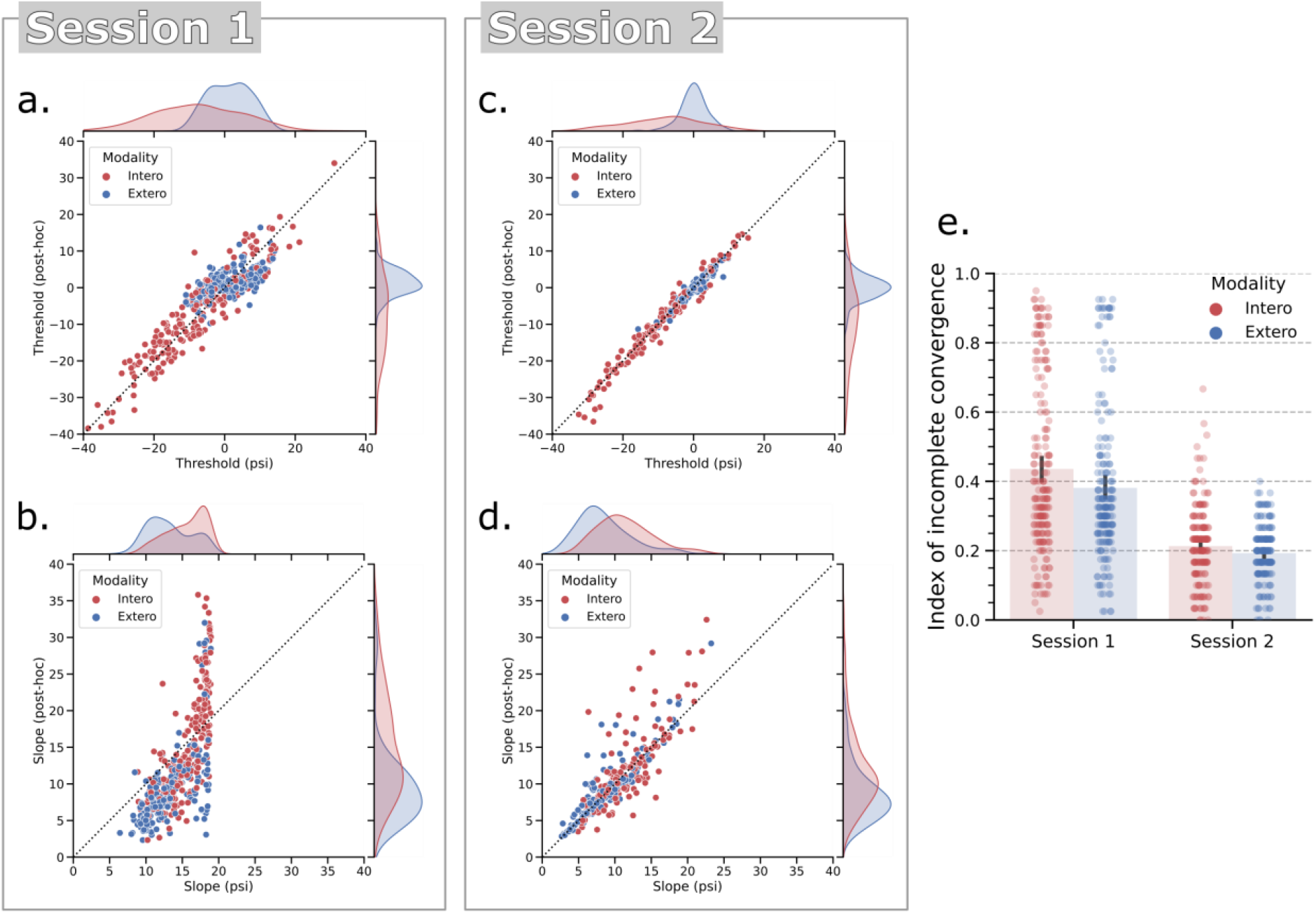
Comparison between online and post hoc Bayesian estimation of slope and threshold parameters of the psychometric functions. Adaptive Bayesian staircases can be biased if their initial parameter settings poorly fit the underlying generative psychometric function, or if a subject makes unrepresentative responses early in the experiment. For example, in this sample we observed that the prior width [0 − 20] on the slope parameter was too low, resulting in a ceiling effect that biased our estimates in a subset of participants. One solution to control these biases is to implement post hoc Bayesian modelling of the observed psychophysical data. We thus re-analyzed the responses for each participant and each condition separately using a Bayesian model to fit a cumulative normal distribution. **A.** The thresholds estimates remained stable, although with a reduced variance for the exteroceptive condition. **B.** The ceiling effect on the slope was normalized by the post hoc modelling, which shifts the posterior mass away from the extremes. The post hoc procedure can thus improve the estimation of the interoceptive and exteroceptive psychophysical parameters. In session 2, both threshold (**C.**) and slope (**D.**) were more reliably estimated after changes we made on the experimental design and prior ranges of the Psi parameters. **E.** We created an index of staircase convergence to quantify the estimation errors observed in session 1 (see below for details). A higher value reflects more imbalanced intensities around the threshold, which is often associated with improper estimates and convergences of the staircases.

Another reason for using a Bayesian model was the presence of incomplete convergence of the Psi staircase during the first session. The Psi algorithm (Kontsevich & Tyler, 1999) is designed to test intensity values that would first increase the precision of the posterior density for threshold. When this confidence around the threshold level is high enough, the staircase starts to improve precision for the slope estimate by testing intensity values around the threshold. This results in a recognizable pattern of higher and lower intensities values alternating regularly around the inferred threshold. Interfering with the Bayesian updating during the first 30 trials of the task, as in Session 1, could result in biased estimation of threshold values. Further, erroneous responses during the first trials may hinder convergence.

Here, we quantified the amount of incomplete Psi staircase convergence through the two sessions. Incomplete convergence is characterized by stimulus intensity values that are consistently higher or lower than the inferred threshold even at the end of the task. To quantify this effect, we calculated an incomplete convergence index using the ratio of high-intensity versus low-intensity values compared to the inferred threshold in the last 40 trials. This ratio was then converted using the following formula:

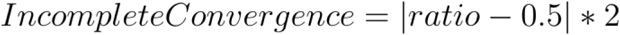

This formula returns a real number between 0 and 1. 0 indicates that the intensity values were equally distributed around the inferred threshold in the last 40 trials. Instead, 1 indicates divergence between the tested intensity values and the inferred threshold. The incomplete convergence indexes for Interoception and Exteroception through Session 1 and 2 are reported in **Supplementary Material Fig. 2. e**). These results revealed a high proportion of incomplete convergence in the first session, in both interoception and exteroception conditions. For example, setting an arbitrary threshold for quality assessment at 0.5 revealed that 63 and 41 participants had poor convergence for interoception and exteroception, respectively. These numbers dropped radically in Session 2 (see **Material and Method**) and corresponded to only 3 and 0 staircases for interoception and exteroception, respectively. The improved convergence in Session 2 is likely due to the introduction of different design choices, aimed at solving the convergence issues observed in Session 1.

### Psychometric results (Session 2)

We reproduced the approach used in the first session and compared threshold and slope values between the interoception and the exteroception conditions. This revealed that during interoception participants had significantly lower psychometric thresholds (mean_Intero_ = −8.50, CI_95%_ [−10.06, −6.92], mean_Extero_ = 0.01, CI_95%_ [−0.47, 0.52], t_(190)_ = −11.15, p < 0.001, BF_10_ = 2.85e+19, d = −1.03) and higher psychometric slopes (mean_Intero_ = 11.96, CI_95%_ [11.22, 12.74], mean_Extero_ = 8.69, CI_95%_ [8.14, 9.28], t_(190)_ = −7.29, p < 0.001, BF_10_ = 9.12e+08, d = 0.67). See Fig. 2 and Supplementary Fig. 1 for illustration of these effects. Similarly to the results in the first session, the negative bias of the threshold parameters suggests that participants underestimated their heart rate on average. The greater slope on the other side, indicates a less precise decision process.

**Legend Supplementary Material 3:**
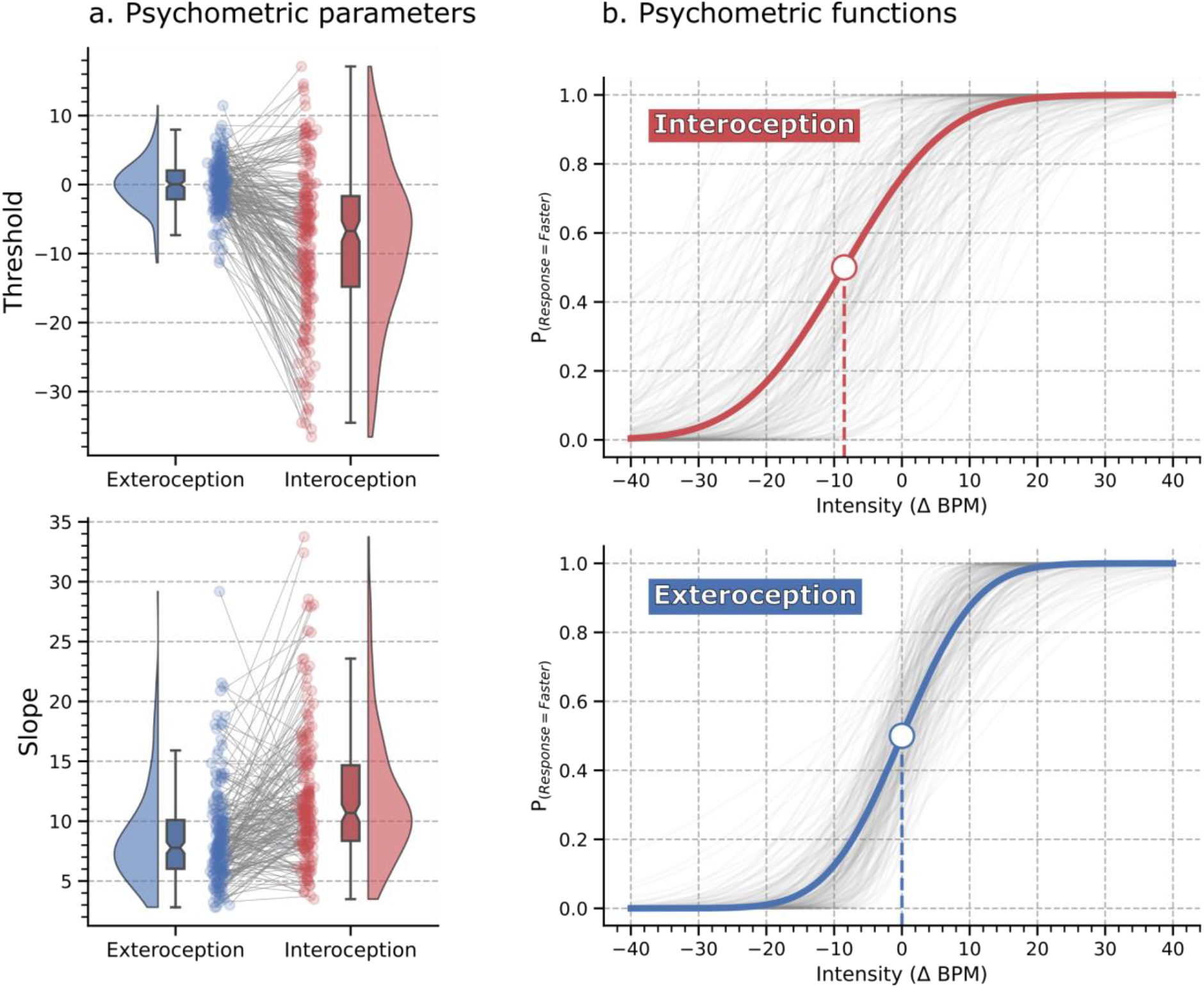
Psychometric parameter estimates and fitted interoception and exteroception psychometric functions (Session 2). **A.** Repeated measures raincloud plots visualizing threshold and slope parameters of the psychometric functions across the two modalities (interoception and exteroception). Data points for every individual are connected by a grey line to highlight the repeated measure effect. **B.** The grey lines show individual subject fits. The dark red and blue lines show the grand mean psychometric function, depicting averaged threshold and slope. Grand mean thresholds are marked by the large point, where the psychometric function crosses 0.5 on the ordinate axis. We observed a strong effect of interoception on both slope and threshold parameters as compared to the exteroceptive control condition.

### Metacognition results (Session 2)

The *d*’, which reflects discimination sensitivity, was lower in the interoception condition (mean_Intero_ = 1.88, CI_95%_ = [1.78, 1.96], mean_Extero_ = 2.25, CI_95%_ = [2.21, 2.3], *t*_(189)_ = −8.10, *p* < 0.001, BF_10_ 9.67e+10, *d* = −0.77). Further, as in the first session, we found that metacognitive sensitivity was significantly lower during interoception. The interoceptive M-ratio estimates were lower (mean_Intero_ = 0.83, CI_95%_ = [0.8, 0.87]) than the exteroceptive ones (mean_Extero_ = 0.96, CI_95%_ = [0.92, 1.01]). The posterior distribution of the repeated measure effect was also lower (mean = −0.17 HDI_94%_ = [−0.28, −0.05].

**Legend Supplementary Material 3:**
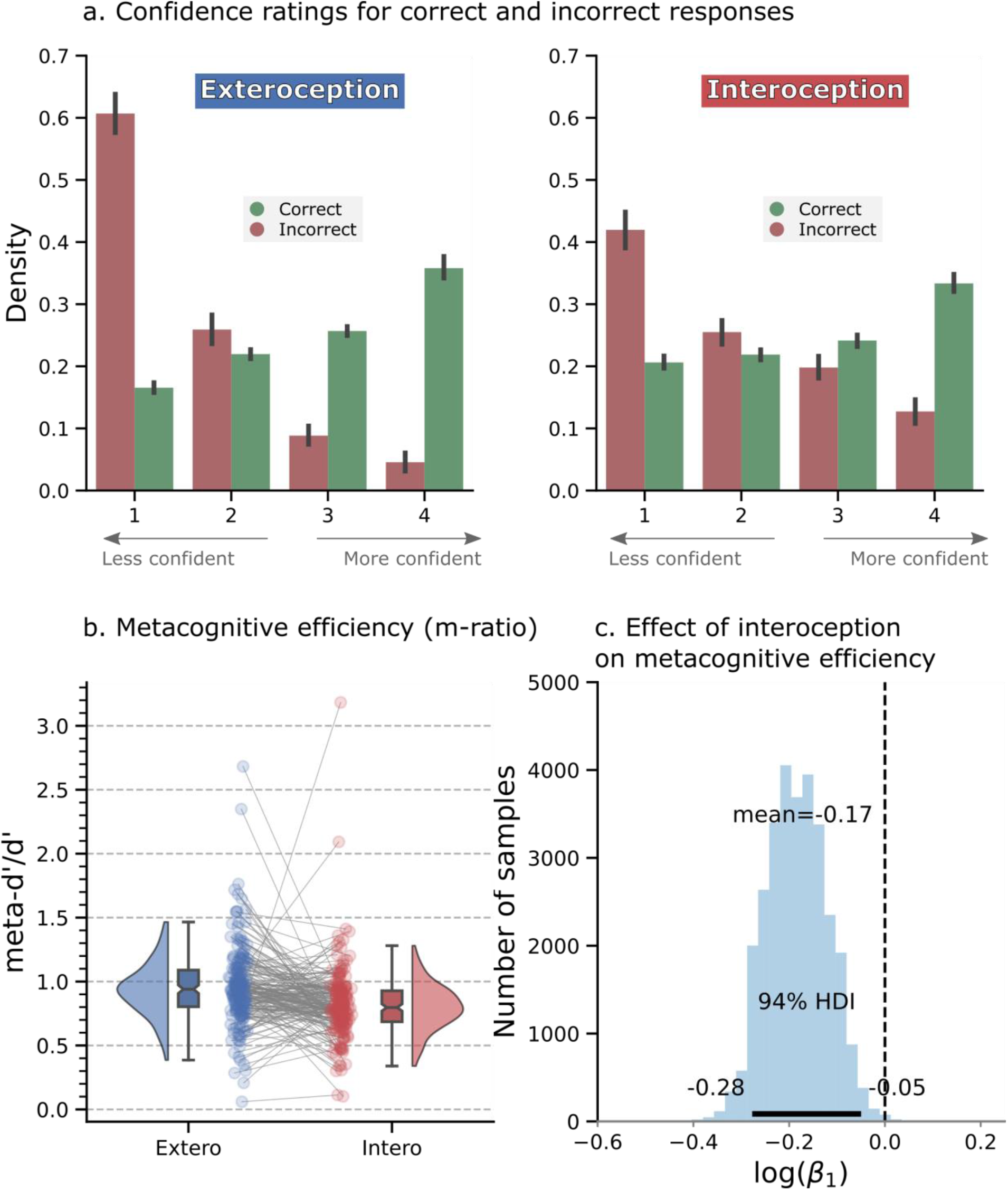
Visualization of metacognitive performance for interoceptive and exteroceptive conditions (Session 2). **A.** Histogram showing the distribution of binned confidence ratings for correct (green) vs. error (red) trials. Higher bins represent higher confidence ratings. Participants were significantly less confident overall in the interoceptive condition and showed reduced calibration as indicated by the flattening of the confidence distributions. To quantify this effect, we estimated “metacognitive efficiency”, a signal theoretic model of introspective accuracy which controls for differences in type 1 (discrimination) performance. Here, an M-ratio of 1 indicates optimal metacognition according to an ideal observer model, whereas values lower than this indicate inefficient use of the available perceptual signal. **B.** This model demonstrated that metacognitive efficiency was substantially decreased for interoceptive relative to exteroceptive judgements. **C.** Histogram of posterior samples from the beta value coding the effect of interoception.

### Cross-modal correlation (Session 2)

We observed more robust and consistent correlations between task parameters within each modality (interoception or exteroception), but few significant correlations between task modalities, indicating a high degree of independence between performance on the two task conditions. Across modalities, response times during the decision process (type 1 measure) were correlated between the interoception and the exteroception conditions (r_s_ = 0.66, CI_95%_ = [0.58, 0.74], *p* < 0.001, n = 190, n_outliers_ = 5), as well as between confidence ratings (r_s_ = 0.54, CI_95%_ = [0.44, 0.64], p < 0.001, n = 190, n_outliers_ = 3).

**Legend Supplementary Material 4:**
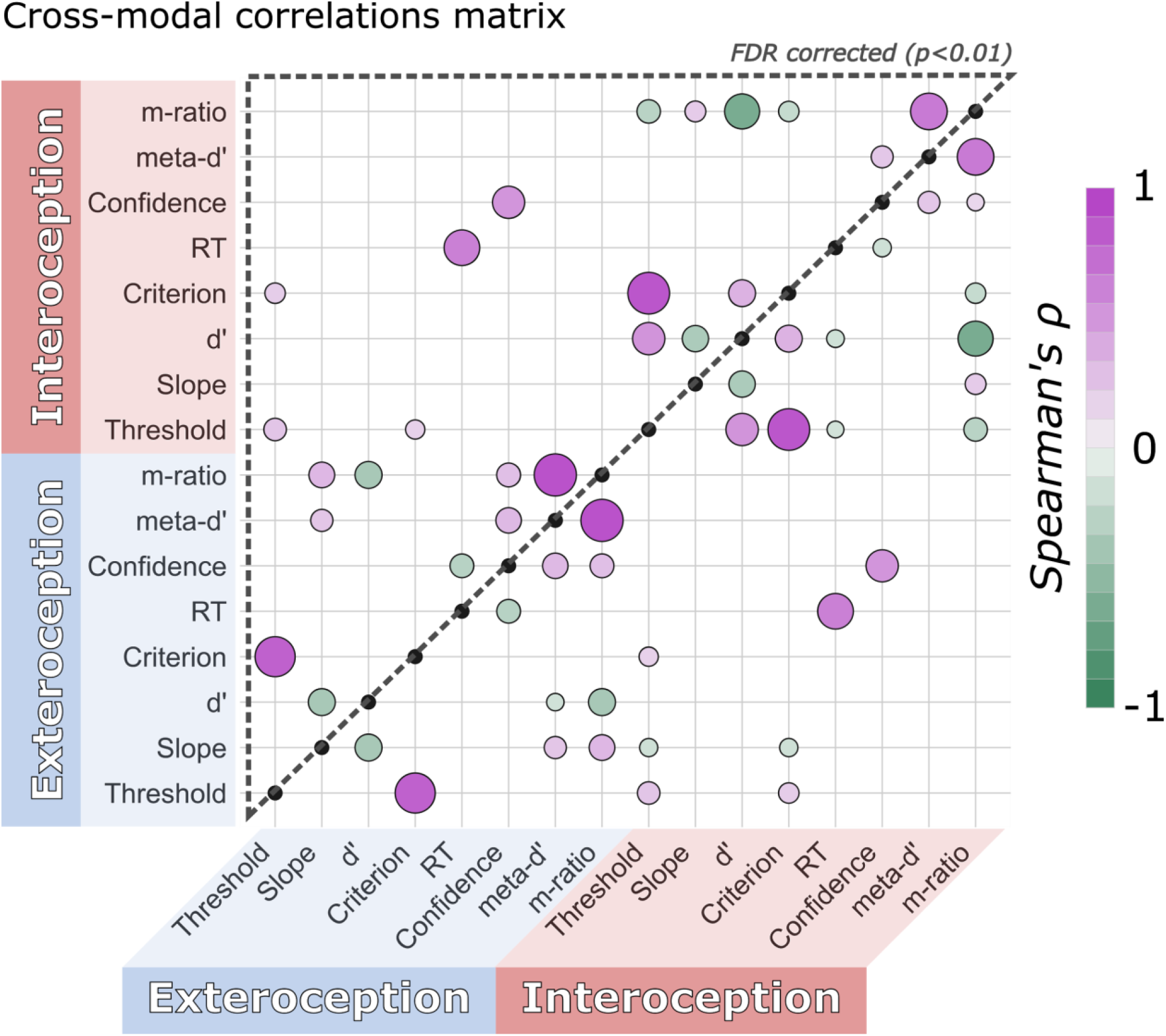
Cross-modal correlation heatmap of task parameters for interoception and exteroception conditions (Session 2). We replicated several of the observations reported in Session 1. Behavioural results were correlated within modalities but with limited dependence across modalities. The only exceptions, already observed in Session 1, were confidence ratings and response times (RT). The figure depicts significant skipped Spearman correlations. **The upper triangle shows results surviving FDR correction (p_FDR_ < 0.01)**. The colour and size of individual points indicate the sign and strength of the estimated correlation coefficients.

